# Metabolic, Fibrotic, and Splicing Pathways Are All Altered in Emery-Dreifuss Muscular Dystrophy Spectrum Patients to Differing Degrees

**DOI:** 10.1101/2022.05.20.492778

**Authors:** Jose I. de las Heras, Vanessa Todorow, Lejla Krečinić-Balić, Stefan Hintze, Rafal Czapiewski, Shaun Webb, Benedikt Schoser, Peter Meinke, Eric C. Schirmer

**Affiliations:** Institute of Cell Biology, University of Edinburgh, Edinburgh, UK; Friedrich-Baur-Institute, Department of Neurology, LMU Clinic, Ludwig-Maximillians-University, Munich, Germany; Wellcome Centre for Cell Biology, University of Edinburgh, Edinburgh, UK

**Keywords:** Emery-Dreifuss, muscular dystrophy, EDMD, skeletal muscle

## Abstract

Emery-Dreifuss muscular dystrophy (EDMD) is a genetically and clinically variable disorder. Previous attempts to use gene expression changes find its pathomechanism were unavailing, so we here engaged a functional pathway analysis. RNA-Seq was performed on cells from 10 patients diagnosed with an EDMD spectrum disease with different mutations in 7 genes. Upon comparing to controls, the pathway analysis revealed that multiple genes involved in fibrosis, metabolism, myogenic signaling, and splicing were affected in all patients. Splice variant analysis revealed alterations of muscle-specific variants for several important muscle genes. Deeper analysis of metabolic pathways revealed a reduction in glycolytic and oxidative metabolism and reduced numbers of mitochondria across a larger set of 14 EDMD patients and 7 controls. Intriguingly, the gene expression signatures segregated the patients into three subgroups whose distinctions could potentially relate to differences in clinical presentation. Finally, differential expression analysis of miRNAs changing in the patients similarly highlighted fibrosis, metabolism, and myogenic signaling pathways. This pathway approach revealed a clear EDMD signature that can both be used as the basis for establishing a biomarker panel specific to EDMD and direct further investigation into its pathomechanism. Furthermore, the segregation of specific gene changes into three distinct categories that appear to correlate with clinical presentation may be developed into prognostic biomarkers, though this will first require their testing in a wider set of patients with more clinical information.

## Introduction

Emery-Dreifuss muscular dystrophy (EDMD) is a genetically heterogeneous neuromuscular orphan spectrum disease affecting ∼0.3-0.4 in 100,000 people ^1^, with clinical variability presenting even in family members carrying the same mutation ^2–5^. EDMD patients present typically in mid to late childhood with early contractures of elbows and Achilles’ tendons and progressive wasting of the lower leg and upper arm muscles. Cardiac involvement is also highly characteristic but tends to appear later in development and quite variably in time, though it tends to be reasonably uniform in the form it takes of cardiac conduction defects and dilated cardiomyopathy ^6^. Other features vary considerably in clinical presentation, leading to the usage of ‘Emery-Dreifuss-like syndromes’ ^7; 8^: patients from the same pedigree can show remarkable phenotypic variation ^2^. The genetic variability is underscored by several confirmed linked genes and several additional candidate genes, although there are still some cases where no confirmed or candidate disease allele has been identified ^9–11^. The lack of large pedigrees in combination with its genetic heterogeneity, clinical variability, already some known modifier genes, and limited patient numbers makes solving its pathomechanism difficult.

The original genes linked to EDMD, *EMD* encoding emerin and *LMNA* encoding lamin A, have both cytoskeletal and gene regulation roles leading to strong arguments for either function being responsible for the EDMD pathomechanism ^12; 13^. The subsequent linking of nesprin and Sun proteins to EDMD ^14; 15^ failed to lend clarity since they function in mechanosignal transduction ^16^. However, several recently linked genes have clear roles in genome organization and regulation ^10^, suggesting this is the pathomechanism. These genes encode proteins that, like emerin, are nuclear envelope transmembrane proteins (NETs) and seem to function by fine-tuning muscle gene expression by promoting the release of pro-myogenic genes from the nuclear periphery to enhance their activation while concomitantly recruiting metabolism genes (many from the alternative differentiation pathway of adipogenesis) to the nuclear envelope to better repress them ^17–19^. EDMD mutations were found in 5 muscle-specific NETs with this genome organization function, PLPP7 (also known as NET39), WFS1, TMEM38A, TMEM201, and Tmem214, and each tested had some specificity in the sets of genes that they target, though there was also some overlap ^17^. These studies together with the wide range of lamin gene regulatory activities led us to the distinct and non-traditional hypothesis for the EDMD pathomechanism where moderate reductions in many genes could have the same phenotype as a shutdown of a single gene on a particular pathway. Accordingly, we considered that searching for uniformity with a pathway analysis might be more revealing than searching for uniformity in the regulation of particular genes.

The only previous study, to our knowledge, using gene expression changes to identify critical misregulated genes underlying EDMD pathophysiology, focused on the identification of genes altered specifically in EDMD compared to a set of 10 other muscular dystrophies ^20^. This study only considered 8 total *LMNA*- or *EMD*-linked cases of EDMD, but EDMD now has many more genes and modifiers linked to it and, moreover, there is a wider clinical spectrum of EDMD-like phenotypes ^11^. Their analysis indicated potential abnormalities in the regulation of cell cycle and myogenic differentiation, associated with perturbations in the pRB/MYOD/LMNA hub, which were consistent with changes in an *Emd^-/-^* mouse model ^21^. Roughly a fifth each of EDMD mutations occurs in *LMNA* and *EMD* while another 5-6% are collectively caused by four other widely expressed nuclear envelope proteins nesprin 1 (encoded by *SYNE1*), nesprin 2 (encoded by *SYNE2*), Sun1 (encoded by *SUN1*) and FHL1 (encoded by *FHL1*) ^14^; ^15^;^22–24^). Another approximately 20% of EDMD mutations were accounted for by muscle-specific NETs that regulate muscle-specific genome organization ^10^. These include NET39 (encoded by *PLPP7*), TMEM38A (encoded by *TMEM38A*), WFS1 (encoded by *WFS1*), NET5 (encoded by *TMEM201*), and TMEM214 (encoded by *TMEM214*) that affect 3D gene positioning with corresponding effects on expression ^17; 19^. Accordingly, we sought to search for commonly affected pathways from a much wider range of EDMD-linked genes including *LMNA*, *EMD*, *FHL1*, *SUN1*, *SYNE1*, *PLPP7*, and *TMEM214* alleles on the expectation that the most important pathways for EDMD pathophysiology would be highlighted.

## Material and methods

### Patient materials

The sources of patient samples were the Muscle Tissue Culture Collection (MTCC) at the Friedrich-Baur-Institute (Department of Neurology, Ludwig-Maximilians-University, Munich, Germany) and the MRC Centre for Neuromuscular Disorders Biobank (CNDB) in London.

### Ethical approval and consent to participate

All materials were obtained with written informed consent of the donor at the CNDB or the MTCC. Ethical approval of the Rare Diseases biological samples biobank for research to facilitate pharmacological, gene and cell therapy trials in neuromuscular disorders is covered by REC reference 06/Q0406/33 with MTA reference CNMDBBL63 CT-2925/CT-1402, and for this particular study was obtained from the West of Scotland Research Ethics Service (WoSRES) with REC reference 15/WS/0069 and IRAS project ID 177946. The study conduct and design complied with the criteria set by the Declaration of Helsinki.

### Myoblast culture and in vitro differentiation into myotubes

Myoblasts were grown in culture at 37 ^O^C and 5% CO_2_ using a ready to use formulation for skeletal muscle (PELOBiotech #PB-MH-272-0090) and maintained in subconfluent conditions. In order to induce differentiation, the cells were grown to confluency and 24 h later the growth medium replaced with skeletal muscle differentiation medium (Cell Applications #151D-250). The differentiation medium was replaced every other day. Myotubes were selectively harvested after 6 days by partial trypsinization followed by gentle centrifugation (Figure S1). Each differentiation experiment was performed in triplicate on different days.

### RNA extraction

Total RNA was extracted from each sample and separated into a high molecular weight fraction (> 200 nt, for mRNA-Seq) and a low molecular weight fraction (< 200 nt, for miRNA-Seq) with the Qiagen RNeasy (#74134) and miRNeasy (#1038703) kits, according to the manufacturer’s instructions. RNA quality was assessed with a Bioanalyzer, and all samples had a RIN > 7, with an average of 9.4 (Table S1).

### mRNA-Seq analysis

Between 3-5ug of total RNA were sent to Admera Health LLC. (NJ, USA) for sequencing in paired-end mode, 2x 150 nucleotides, using an Illumina HiSeq 2500 sequencer. The sequencing library was prepared with the NEBNext Ultra II kit, with RiboZero rRNA depletion (NEB #E7103). Between 60-90 million paired end reads were obtained from each sample and mapped to the human genome (Hg38) with STAR v2.7.5a ^96^ using default parameters. Mapping quality was assessed with FastQC v0.11.9 (https://www.bioinformatics.babraham.ac.uk/projects/fastqc). Sequencing adaptors were removed with trimmomatic v0.35 ^97^. Low quality reads and mitochondrial contaminants were removed, leaving on average 70 million useful reads per sample (Table S1). Differential expression analysis was performed in R with DESeq2 v1.32.0 ^98^ after transcript quantitation with Salmon v1.4.0 ^99^. We used an FDR threshold of 5% for differential expression.

### miRNA-Seq analysis

miRNA was sent to RealSeq Biosciences Inc. (SC, USA) for sequencing using an Illumina NextSeq 500 v2 sequencer in single end mode, 1x 75 nucleotides. The sequencing library was prepared with Somagenics’ Low-bias RealSeq-AC miRNA library kit (#500-00012) and quality assessed by Tapestation (Lab901/Agilent). On average 5 million good quality reads were obtained per sample. Mapping and quality trimming was performed using the NextFlow nf-core/smrnaseq pipeline (web resources) with default parameters, which summarizes the reads per miRNA using the annotations from mirTop (web resources). Differential expression analysis was performed in R with DESeq2 v1.32.0. We used an FDR threshold of 0.2 for differential expression. Putative miRNA targets were extracted from miRDB (web resources) for each differentially expressed miRNA and their expression compared against the miRNA. We kept as potential targets those genes whose expression changed in the opposite direction of the miRNA.

### Bakay muscular dystrophy dataset analysis

Normalized (MAS5.0) microarray transcriptome data for a panel of 11 muscular dystrophies and healthy controls were downloaded from the Gene Expression Omnibus database (GEO), accession GSE3307 (web resources). Differential expression analysis comparing each disease to the controls was performed using Limma 3.48.1 ^100^. We used an FDR threshold of 5% for differential expression.

### Functional analyses

Functional analyses were performed with g:Profiler ^101^ and Gene Set Enrichment Analysis (GSEA v4.1.0) ^35^ tools. g:Profiler was used to determine enriched categories within a set of DE genes, with an FDR of 5% as threshold. GSEA was performed with default parameters, in particular using ‘Signal2Noise’ as ranking metric and ‘meandiv’ normalization mode. Redundancy in category lists was reduced by comparing the similarity between each pair of enriched categories using Jaccard similarity coefficients. Hierarchical clustering (k-means) was then applied to the resulting matrix in order to identify groups of similar functional categories, and a representative from each group chosen. Full unfiltered results are shown in Supplemental Table S3. Tissue-specific gene enrichment analysis was evaluated with TissueEnrich ^102^.

For the miRNA-Seq experiments, functional analysis was first performed using g:profiler on the set of DE miRNA genes. Then, putative targets for each miRNA were extracted and their expression compared to the relevant miRNA. Putative targets whose expression was not altered in the opposite direction as the miRNA were removed from the list. Significant functions were displayed using Cytoscape v3.8.2 ^103^, with the size of the functional labels proportional to the number of miRNAs assigned to each function.

### Real time metabolic measurements

Metabolic measurements on primary human myoblast cultures were performed using a Seahorse XFp Extracellular Flux Analyzer (*Agilent Technologies*). For this, myoblasts of matched passage number were seeded in XFp Cell Culture Miniplates (103025-100, *Agilent Technologies*) at a density of 1.5×10^4^ cells per well. Cell density was assessed using an automated cell counter (TC20, BioRad). Oxygen consumption rates (OCR) respectively extracellular acidification rates (ECAR) were measured using the Mito Stress Test Kit and the Glycolysis Stress Test Kit (*Agilent Technologies*) according to the manufacturer’s instructions. Samples were measured in triplicates and each measurement was repeated between two and four times. Data were normalized to the number of cells and analyzed for each well.

### Fuel dependency tests

Glucose dependency and fatty acid dependency were determined according the instruction of Agilent Seahorse XF Mito Fuel Flex Test kit (Agilent). The glutamine dependency was determined from the glucose and fatty acid measurements.

### Mitochondrial gene quantification

Reverse transcription of RNA was performed using the QuantiTecT Reverse Transcription Kit (Qiagen) following the manufacturer’s instructions. For the reaction we used the SYBR® Green Master Mix (Bio-Rad) and samples were run and measured on CFX Connect^TM^ (Bio-Rad). As genome reference gene *B2M* (FP: 5‘-TGCTGTCTCCATGTTTGATGTATCT-3‘; RP: 5‘-TCTCTGCTCCCCACCTCTAAGT-3‘) ^104^ Primer sequences for the mitochondrial genome were: FP: 5′-TTAACTCCACCATTAGCACC-3′; RP: 5′-GAGGATGGTGGTCAAGGGA-3′.^105^ Samples were analyzed using the delta Ct method.

### Splice site prediction analysis

Raw data was mapped to the human genome assembly GRCh38 (hg38) and sorted by coordinate using STAR 2.7.9a ^96^ for analysis in DESeq2 ^98^ and DEXSeq ^106^, trimmed using an in-built trimming function for rMATS ^107^ or counted using Kallisto 0.48.0 ^108^ for isoformSwitchAnalyzer (ISA) ^109^. All analyses were performed for G1, gp1, gp2, and gp3 separately. Visualizations were conducted in R version 4.1.2.

DESeq2: Mapped reads were counted using FeatureCounts, then analysed using DESeq2 ^98^. The R package fgsea was used for gene set enrichment analysis, genes were assigned to biological pathways retrieved from MSigDB v7.5.1 (c5.go.bp.v7.5.1.symbols.gmt, 7658 gene sets, ^35^). Splicing pathways and their genes were plotted using GOPlot ^110^.

DEXseq: Mapped reads were counted using the in-built DEXseq counting function in python 3.9. Standard DEXseq workflow was followed. Exons with logFCs > |1| and p-values < 0.05 were set to be significantly different.

rMATS: Standard workflow was followed, code was executed in python 2.7. Results were analysed in R and set to be significantly differentially spliced with psi-values > |0.1| and p-values < 0.05. Pie charts displaying the distibution of event usage were generated for all groups (Supps. something) GO term enrichment analyses was performed using g:profiler2^111^. All events were searched for muscle specific genes using a set of 867 genes relevant for muscle system process, development, structure and contraction, combined from GO terms (GO:0003012, GO:0006936, GO:0055001 and GO:0061061).

ISA: Kallisto counts were read into R and standard ISA procedure was followed, including Splicing analysis using DEXseq, coding potential using CPC 2.0 ^112^, domain annotation using HmmerWeb Pfam 35.0 ^113^, signal peptides using SignalP 5.0 ^114^ and predicition of intrisically unstructured proteins using IUPred2A ^115^. In-built visualization tools were used for splicing maps.

## Results

### RNA-Seq Analysis of EDMD Patient Cells

We performed RNA-Seq on myotubes differentiated *in vitro* from myoblasts isolated from 10 unrelated EDMD patients with distinct mutations in 7 different genes to sample the genetic diversity of EDMD (**Figure 1**). Patient mutations were TMEM214 p.R179H, PLPP7/NET39 p.M92K, Sun1 p.G68D/G388S, Nesprin 1 p.S6869*, Emerin p.S58Sfs*1, FHL1 mutations c.688+1G>A, p.C224W, and p.V280M, and Lamin A mutations p.T528K and p.R571S. These mutations covered a wide range of clinical phenotypes with the age of onset ranging from early childhood to adult life and associated pathology ranging from no reported contractures to rigid spine (**Table 1**). Myoblast isolation followed by *in vitro* differentiation was chosen over directly isolating mRNA from the tissue samples in order to try to capture the earliest changes in gene expression due to the disease mutations and to reduce tertiary effects and variation from the age of the patients, range in time from onset to when biopsies were taken, and differences in biopsy site (**Table 1**). These patient variables could also affect the efficiency of myotube differentiation; so, to ensure that different percentages of undifferentiated cells in the population did not impact measuring gene expression changes, the myotubes were specifically isolated by short trypsinization thus removing all myoblast contamination (**Figures 1** **and S1**). To define the baseline for comparison, 2 age-matched healthy controls were similarly analyzed. Samples from all patients and controls all yielded high-quality reads ranging between 28 and 47 million paired-end reads (**Table S1**).

**Figure 1.**
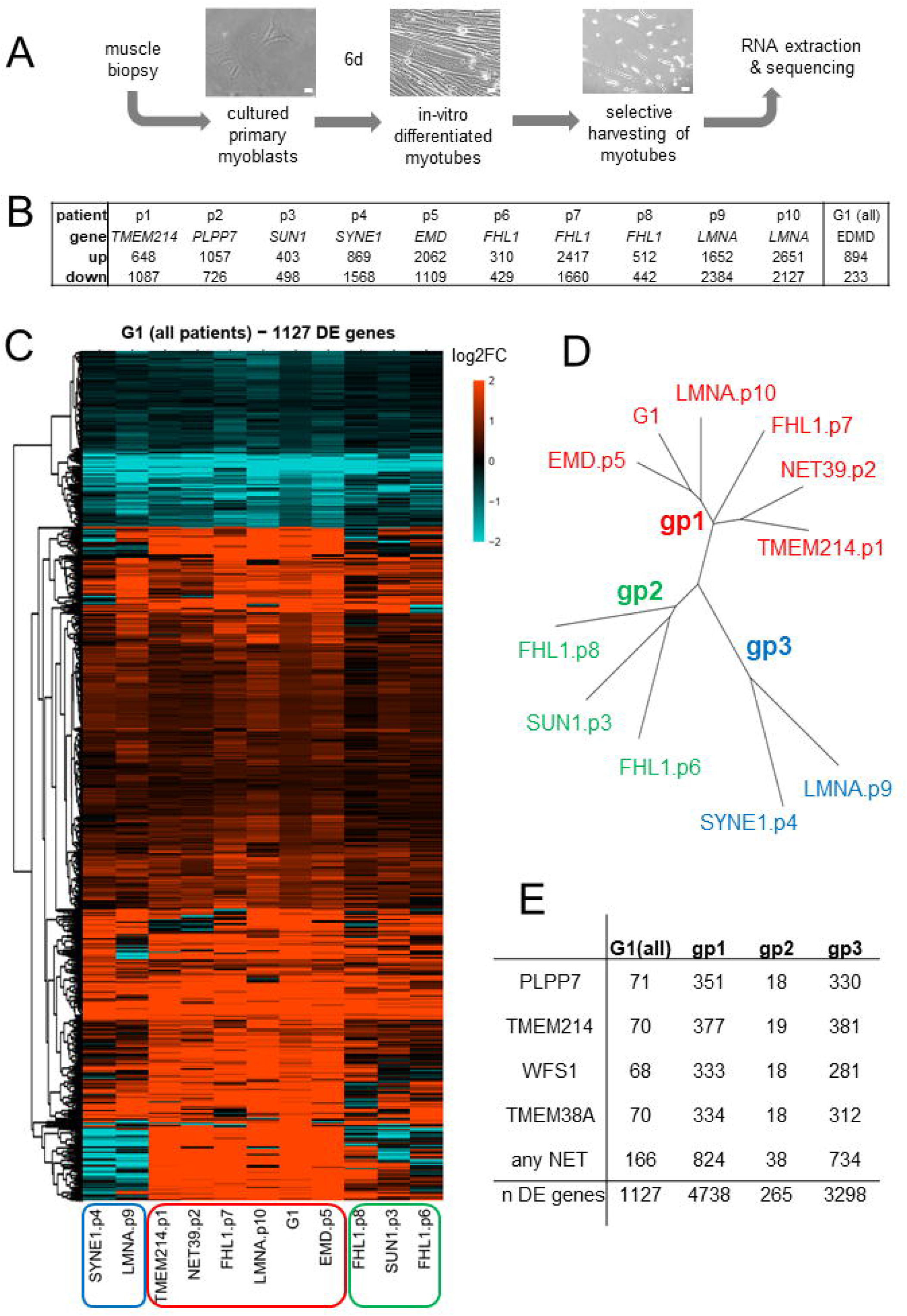
RNA-Seq for EDMD. A. Workflow. Muscle biopsies were taken from the regions described in Table 1 and myoblasts recovered. These were then differentiated *in vitro* into myotubes, and myotubes selectively recovered by partial trypsinization. Myotube RNA was extracted and used for sequencing. Scale bar 20µm. B. Number of genes differentially expressed (FDR 5%) for each individual patient and for all patients considered together as a single group (denoted as “G1”). C. Heatmap of log2FC values for genes changing expression in EDMD patients compared to healthy controls (G1). Red is upregulated and blue is downregulated. Black indicates no change. The G1 lane is the averaged data across all patients. D. Dendrogram showing the relationships between patients, which fall into three broad groups. E. Overlaps with genome-organizing NETs gene targets. The numbers of genes altered by knockdown of muscle-specific genome-organizing NETs Wfs1, Tmem38a, NET39, or Tmem214 that were also altered in the EDMD patient cells is given. G1 refers to the analysis of all 10 patients as a single group against the healthy controls, while gp1-3 refers to the three subgroups identified. Total number of differentially expressed (DE) genes are given for each.

**Table 1.** Patients and Controls used in this study. List of patients and controls including gene, mutation, age at biopsy, age of onset, sex, creatine kinase (CK) levels, clinical features, and the muscle groups from whence biopsies were obtained. Note that the normal range for CK levels is 25-200 U/l.

| ID | gene | mutation | RNAseq | metabolic analysis | age | age of onset | sex | CK(U/l) | muscle weakness | contractures | cardiac defects | neurological defects | biopsy location | other clinical information |
| --- | --- | --- | --- | --- | --- | --- | --- | --- | --- | --- | --- | --- | --- | --- |
| C1 | control |  | yes | yes | 35 |  | M |  | no | no | no |  | Gastrocnemius caput laterale |  |
| C2 | control |  | yes | yes | 13 |  | M |  | no | no | no |  |  |  |
| C3 | control |  |  | yes | 43 |  | M |  | no | no | no |  | Biceps brachii |  |
| C4 | control |  |  | yes | 36 |  | F |  | no | no | no |  | Biceps brachii |  |
| C5 | control |  |  | yes | 49 |  | F |  | no | no | no |  | Vastus lateralis |  |
| C6 | control |  |  | yes | 49 |  | F |  | no | no | no |  |  |  |
| C7 | control |  |  | yes | 53 |  | M |  | no | no | no |  | Vastus lateralis |  |
| P1 | TMEM214 | p.179H | yes | yes | 48 | 28 | F | 350 | yes |  | no |  | Rectus femoris | LGMD, ptosis, type 2 fiber atrophy |
| P2 | PLPP7 | p.M92K | yes | yes | 34 | 16 | F | 400 | yes |  | no |  | Vastus lateralis | distal myopathy |
| P3 | SUN1 | p.G68D<br>p.G388S | yes | yes | 9 | 10 | M | >3000 | yes | yes | yes |  | Quadriceps femoris |  |
| P4 | SYNE1 | p.6869*<br>p.6869* | yes | yes | 15 |  | F | 347 | yes |  |  | yes | Quadriceps femoris | growth deficit, ataxia |
| P5 | EMD | p.S58Sfs*1 | yes | yes | 17 | 12 | F | 278 | yes | yes | yes |  | Biceps brachii |  |
| P6 | FHL1 | c.688+1G>A | yes |  | 28 |  | M | 1000 | yes |  | yes |  | Biceps brachii |  |
| P7 | FHL1 | p.C224W | yes | yes | 46 |  | M | 300-1000 | yes |  |  |  | Tibialis anterior |  |
| P8 | FHL1 | p.V280M | yes | yes | 21 |  | M | 231 | yes | yes |  |  | Gastrocnemius caput laterale | rigid spine |
| P9 | LMNA | p.T528K | yes |  | 2 |  | M |  | yes |  |  |  |  |  |
| P10 | LMNA | p.R571S | yes | yes | 24 | 9 | F | up to 12000 | yes |  |  |  |  | degenerative myopathy |
| P11 | LMNA | p.R545C |  | yes | 18 |  | M |  | yes | yes | yes |  |  |  |
| P12 | LMNA | p.R543W |  | yes | 12 |  | F |  | yes |  |  |  | Vastus lateralis | acrogeria |
| P13 | unknown | unknown |  | yes | 14 |  | M | 2800 | yes | yes | yes |  | Tibialis anterior | degenerative myopathy |
| P14 | FHL1 | p.C224W |  | yes | 42 |  | M | 1700 | yes | yes | yes |  | Vastus lateralis | myofibrillar myopathy |

First, we compared each individual patient against the controls. Compared to the controls, each individual patient had between 310 and 2,651 upregulated genes and between 429 and 2,384 downregulated genes with an FDR of 5% (**Figure 1**). The large difference in the number of differentially expressed genes between patients suggested large heterogeneity. When we calculated the intersection of DE genes in all patients, only three genes were similarly downregulated (*MTCO1P12*, *HLA-H*, *HLA-C)*, and one upregulated (*MYH14)* at 5% FDR, indicating a high degree of variation between patients (**Figure S2**). *MTCO1P12* is a mitochondrially-encoded pseudogene that has been reported to be severely downregulated in inflammatory bowel disease, associated with reduced mitochondrial energy production ^25^. HLA-C is a member of the MHC class I and is involved in interferon gamma signaling while HLA-H is a pseudogene derived from HLA-A which may function in autophagy ^26–28^. Mutations in non-muscle myosin gene *MYH14* appear to be associated with hearing loss rather than muscle defects ^29; 30^, although it has also been recently linked to mitochondrial fission defects ^31^.

In order to identify common features despite the above-mentioned heterogeneity, we next compared the patients as a single group against the controls instead of one-by-one and we denoted this comparison as G1 (one single group) throughout the text (**Figure 1**). The G1 comparison revealed a signature of 1127 DE genes (894 upregulated and 233 downregulated). Approximately 60% of these genes identified were altered in the same manner in all 10 patients, although only 4 of them had an associated FDR value below 0.05 (**Figure S2**). The remaining 40% appear to be quite variable between patients in level of expression and/or direction of change. Such distinction in gene expression signatures might also contribute to clinical variation in EDMD.

Hierarchical clustering identified three broad patient subgroups based on gene expression patterns (**Figure 1**). These groupings were: Group 1 - Emerin p.S58Sfs*1, Tmem214 p.R179H, NET39 p.M92K, Lamin A p.R571S, FHL1 p.C224W; Group 2 - Sun1 p.G68D/G388S, Fhl1 c.688+1G>A, Fhl1 p.V280M; Group 3 - Nesprin 1 p.S6869*, Lamin A p.T528K (**Fig. 1D**). The same groups were independently identified using a principal component analysis (PCA) (**Figure S2**). Of particular note, PCA and t-distributed stochastic neighbor embedding (t-SNE) analyses both revealed that clustering was independent of parameters such as patient gender or age or myotube enrichment differences (**Figure S3**). Moreover, the several FHL1 and lamin A mutations tested segregated into different expression subgroups. At the same time, the more recently identified EDMD mutations in TMEM214 and NET39 segregated with more classic emerin, FHL1, and lamin A mutations, further indicating the likelihood that their genome organizing functions could mediate core EDMD pathophysiology.

The hypothesis that EDMD is a disease of genome organization misregulation is underscored by the fact that 15% of the genes changing expression in EDMD patient cells were altered by knockdown of at least 1 of the 4 muscle-specific genome-organizing NETs that we previously tested ^17^ (**Figure 1**). Interestingly, in that study most of the genes altered by knockdown of NET39, TMEM38A, and WFS1 were non-overlapping, while those altered by knockdown of TMEM214 exhibited considerable overlap with the sets altered by each of the other NETs ^17^. However, here there were roughly 70 DE genes overlapping with the sets of genes altered by knockdown of each individual NET while the total number of DE genes under the regulation of any of the four NETs was 165, indicating an enrichment in the EDMD DE set for genes influenced by multiple NETs (**Figure 1**). Thus, it is not surprising that NET39 and TMEM214 were both segregated together. Another interesting observation is that the number of NET-regulated genes overlapping with Group 1 and Group 3 was similar, but much fewer were overlapping for Group 2. This suggests that gene misregulation in Groups 1 and 3 might be more strongly mediated by the muscle-specific NET-gene tethering complexes than in Group 2.

### Functional Pathway Analysis of Gene Expression Changes in EDMD Patient Cells

The primary aims of this study were to determine whether a functional pathway analysis would be more effective at revealing the underlying EDMD pathomechanism than just looking for uniformly altered genes and to identify possible biomarkers in the gene expression signatures. Before using this approach with the wider set of EDMD alleles, we applied a pathway analysis to the data from the previous microarray study by Bakay and colleagues where just *LMNA* and *EMD* mutations were considered ^20^. We reanalyzed Bakay’s EDMD data and extracted the subset of DE genes with FDR of 5% (1349 and 1452 up- and down-regulated genes, respectively). In order to identify enriched functional categories within each set of DE genes, we used g:Profiler ^32^. This tool calculates the expected number of genes to be identified for any given functional category by chance and compares it to the number of genes observed. We selected categories that were significantly enriched with an FDR of 5%. The resulting list was then summarized by selecting representative classes using a similar approach to Revigo ^33^, but extended to other functional category databases in addition to gene ontology (GO) terms. Briefly, similarity matrices were generated by calculating pairwise Jaccard similarity indices between categories and used this information to group together similar functional categories based on the genes identified. Redundancy was then reduced by choosing a representative category from each group.

The functional categories enriched in the set of EDMD-upregulated genes revealed defects in cytokine signaling, organization of the extracellular matrix (ECM), and various signaling pathways important for muscle differentiation and function (*e.g*: PI3K-Akt, TGF-beta, SMADs). In addition, there was an aberrant upregulation of alternative differentiation pathways, notably adipogenesis but also angiogenesis and osteogenesis. The functions highlighted among the downregulated genes were largely related to metabolism, mitochondrial especially, as well as ribosome biogenesis, muscle contraction, and myofibril assembly (**Figure 2**). Applying the same methodology to our wider set of patient alleles highlighted fewer pathways than what we observed in the Bakay EDMD data. Among the upregulated categories, neurogenesis and ECM-related functions stood out, as well as MAPK signaling, lipid transport, and TAP binding which is linked to interferon-gamma signaling ^34^. One category stands out among the downregulated genes: RNA splicing (**Figure 2**). The data above used g:Profiler which is very sensitive to the number of DE genes identified because it looks for statistical overrepresentation of genes belonging to specific functional categories among a set of previously identified DE genes. By contrast, gene set enrichment analysis (GSEA) ^35^ does not prefilter the data and instead ranks all genes according to the difference in expression between the two conditions tested: controls and EDMD. Next, it determines whether the distribution in the ranked list for any given functional category is random or significantly enriched statistically at either end of the ranked list. This method is especially sensitive for detecting functional categories where many genes are altered by a small amount and does not consider individual gene p-values. Therefore, we also applied GSEA to our data, querying several functional genesets within the Reactome, KEGG and WikiPathways databases, found in the Molecular Signatures Database (MSigDB) ^36^. This approach identified a larger set of functional categories that generally expanded on those identified by g:Profiler and matched better what we observed from the Bakay EDMD geneset, with strong links to ECM organization that may be relevant to fibrosis, cytokine signaling, metabolism, differentiation, and splicing (**Figure 2**).

**Figure 2.**
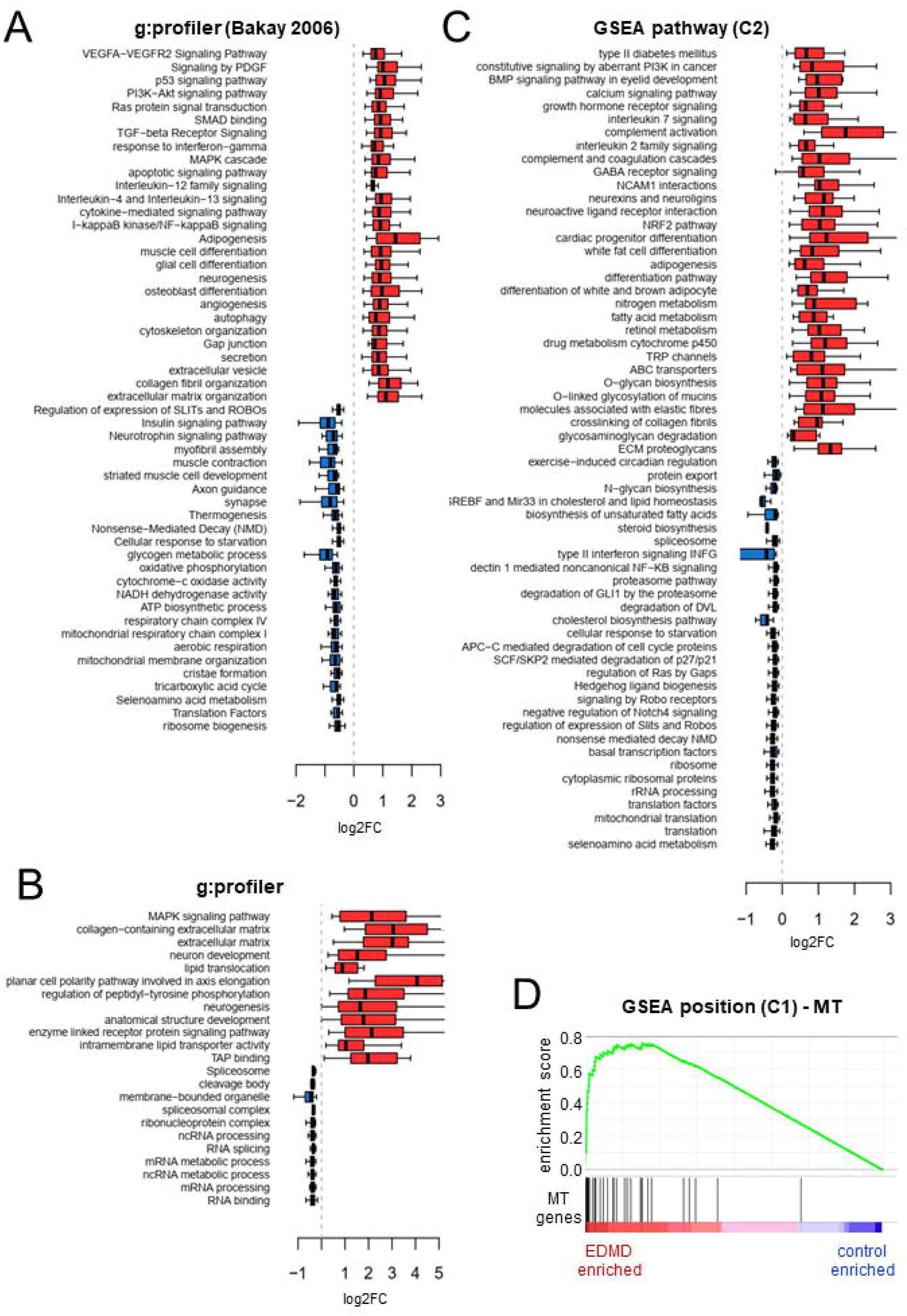
Search for functional categories changing in EDMD patients. A. Box plot of log2FC values for differentially expressed genes within significantly enriched functional categories in EDMD patients analyzed in the Bakay study using g:profiler. B. Box plot of log2FC values for differentially expressed genes within significantly enriched functional categories in EDMD patients analyzed in this study using g:profiler. C. Box plot of log2FC values for leading edge genes within significantly enriched functional categories in EDMD patients analyzed in this study using GSEA pathway (C2 geneset collection: canonical pathways). D. GSEA enrichment plot for mitochondrially encoded genes. GSEA analysis using the C1 geneset (positional) revealed an upregulation of mitochondrially encoded genes. See Supplemental Table S3 for further details.

An expansion of categories for metabolic functions included specific categories for diabetes mellitus, adipogenesis, white and brown fat differentiation, nitrogen metabolism fatty acid metabolism, retinol metabolism, and many others. Similarly, there was an expansion of cytokines supporting inflammation for the fibrotic pathways and proteoglycans and elastin adding to the previous emphasis on collagens for ECM defects. Among the differences between our data and Bakay’s EDMD data, two categories stand out: RNA splicing and calcium signaling, which were only observed in our data. It is unclear how much this reflects using terminally differentiated muscle material versus early stages of differentiation *in vitro*, or a factor of microarray versus RNAseq analysis. In some cases, this is most likely due to the different transcriptome platform used. For example, applying GSEA to genomic positional genesets revealed near uniform upregulation of all mitochondrially encoded genes (**Figures 2** and **3**). This could not be observed on Bakay’s data because the microarrays did not contain probes for mitochondrially encoded genes. The upregulation of mitochondrial transcripts could lead to increased oxidative stress ^37^. This finding provides yet another mechanism that could lead to metabolic dysregulation on top of the alterations already indicated by the nuclear genome transcript changes. This further underscored the need to test for actual metabolic deficits in the patient cells themselves as well as to further investigate the other functional pathways highlighted by this analysis.

**Figure 3.**
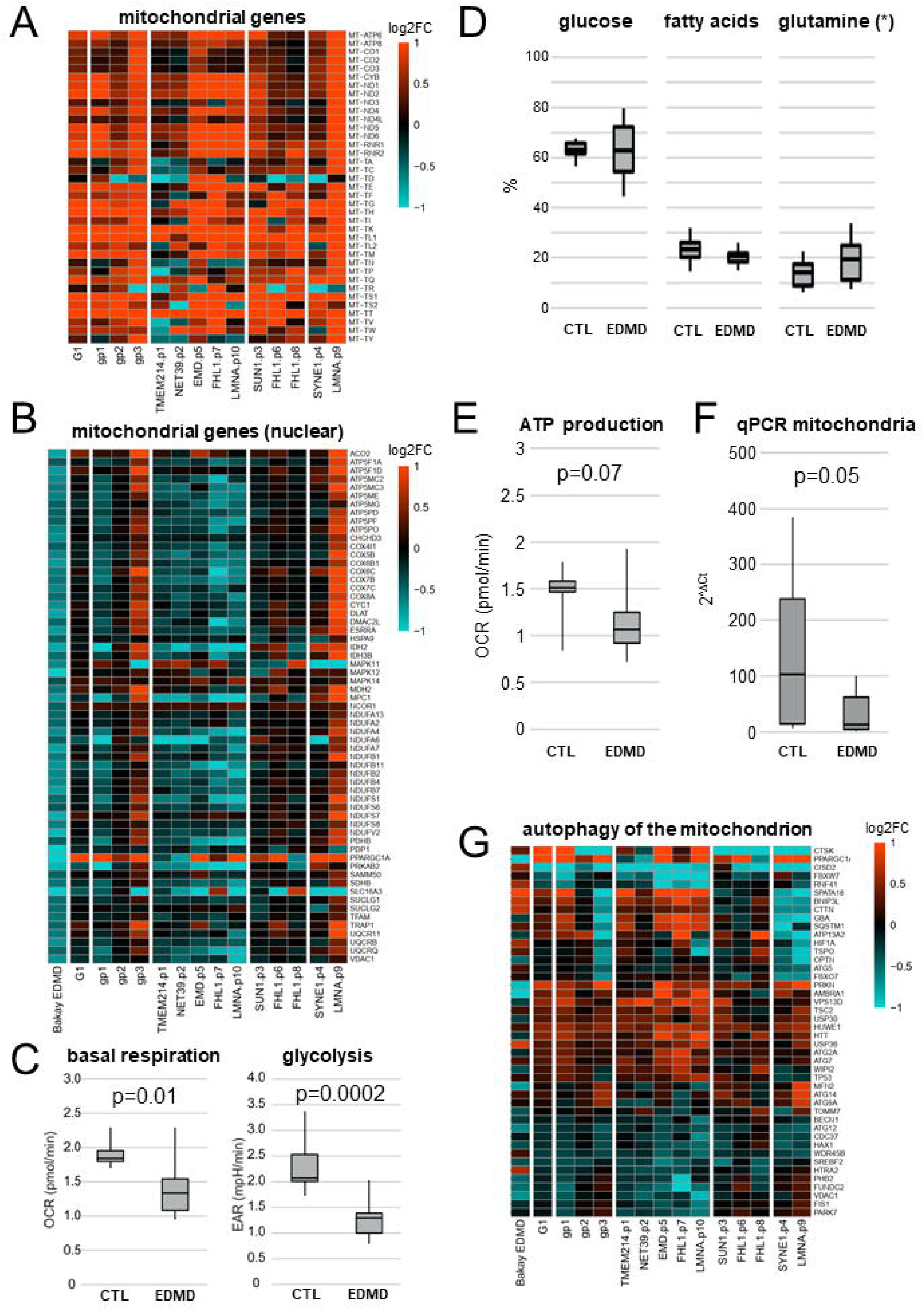
Metabolism changes confirmed in patients. A. Heatmap of all mitochondria-encoded genes showing almost uniform upregulation in EDMD patients. B. Heatmap of nuclear-encoded downregulated genes in Bakay’s study that supported mitochondria function. Downregulation is also generally observed in patients belonging to Group 1, but not Groups 2 or 3. Functional analyses performed in primary patient (n=12) and control (n=7) myoblast cultures for C. glycolysis and basal respiration show a significant reduction of both in EDMD myoblast cultures. D. Testing for fuel dependency we could not observe any changes (* glucose and fatty acids have been measured; glutamine calculated). E. ATP production is reduced in EDMD samples. All functional experiments have been repeated in at least three independent experiments. F. qPCR of mitochondria shows a reduction of mitochondria in EDMD myoblasts. G. Heatmap of mitophagy genes (GO-BP, GO:0000422, Autophagy of the mitochondrion) that are differentially expressed in at least one of Bakay’s EDMD, G1 analysis (all 10 patients analyzed as a single group), or G3 analysis (each of the three subgroups analyzed separately).

### Detailed Analysis of Metabolic Pathways Uniformly Altered in EDMD Patients

Since metabolic disruption has been previously reported to affect muscle differentiation/myoblast fusion ^38^, we decided to investigate this further. While we identified a general upregulation of mitochondrially-encoded genes, Bakay’s data showed a downregulation of several classes related to mitochondrial function (**Figure 2**) which was due entirely to nuclear-encoded genes, as there were no mitochondrial genes represented in the microarrays (**Figure 3**). When we checked the behavior of those genes in our data, we did not observe the same downregulation when considering all 10 patients as a single group (**Figure 3**). However, this is largely due to variability among the patient subgroups identified earlier, suggesting a mechanistic breakdown between them. Group 1 which contained half of our patients, including emerin and lamin A mutations, exhibited the same general downregulation of the nuclear-encoded mitochondrial genes. In contrast, Group 2 displayed no alteration in gene expression, while Group 3 showed upregulation although this was driven mostly by patient 9 (*LMNA*) with the other patient in the group, patient 4 (*SYNE1*), displaying very few changes. While no single gene was uniformly altered in the same direction for all patients, several genes from glycolytic and oxidative metabolism pathways, typically encoding components of mitochondrial complexes, were altered in all tested patients. Other non-mitochondrial metabolic pathways were also altered such as lipid translocation (**Figure 2** **and Table S3**). Interestingly, downregulation of nuclear-encoded mitochondrial genes was also generally observed in other muscular dystrophies included in the study by Bakay and colleagues (**Figure S4**)

To investigate the relevance of these gene changes to cellular metabolism, we performed real-time metabolic analysis using the Seahorse XFp Extracellular Flux Analyzer. Myoblasts isolated from the above patients plus several additional EDMD patients and controls were tested, so that we had a total of 14 EDMD patients and 8 controls for this analysis (**Table 1**). Probing for glycolysis, a significant reduction of the extracellular acidification rate in the EDMD samples was observed (**Figure 3**). Next, we investigated mitochondrial function. When testing for basal respiration there was also a significant reduction of the oxygen consumption rate in the EDMD samples (**Figure 3**). There were no significant differences in fuel dependency, but ATP production was considerably reduced in the EDMD samples (**Figure 3**). The significant reduction in mitochondrial respiration raised another possibility to investigate, that the absolute number of mitochondria might also be down due to problems in mitochondria biogenesis. Therefore, we quantified relative mitochondria numbers using by qPCR. This revealed a clear reduction in mitochondria numbers (**Figure 3**), which with the generally elevated mitochondrial genome transcripts would suggest that a reduction in mitochondria numbers resulted in an over-compensation of expression which in turn could have resulted in inhibiting mitochondrial fission and repair. Thus, we also investigated whether genes in pathways associated with mitophagy were altered in the patients. Indeed, multiple mitophagy pathway genes were altered in all patients (**Figure 3**). Although no one individual gene was altered in all the patients, it is worth noting *CISD2* is significantly downregulated in most patients. Reduction of CISD2 has been linked to degeneration of skeletal muscles, misregulated Ca^2+^ homeostasis and abnormalities in mitochondrial morphology in mouse ^39^, as well as cardiac dysfunction in humans ^40^.

### Detailed Analysis of Other Pathways Uniformly Altered in EDMD Patients

Several studies suggest that the timing of several aspects of myotube fusion could underlie some of the aberrancies observed in patient muscle ^41^ and, though it is unclear whether fibrosis drives the pathology or is a consequence of the pathology, fibrosis has been generally observed in EDMD patient biopsies. Contributing to these processes could be several subpathways that fall variously under the larger pathways for ECM/ fibrosis, cell cycle regulation, and signaling/ differentiation (**Figure 4**). As for the metabolic analysis, no individual genes were altered in cells from all patients, but every patient had some genes altered that could affect ECM through changes in collagen deposition (**Figure 4**). For example, 35 out of 46 collagen genes exhibited changes in at least one comparison (Bakay EDMD, G1 or one of the subgroups gp1, gp2 and gp3) and all patients had multiple of these genes altered (**Figure S5**). Note that it often appears visually that the Bakay data in the first column has little change when viewing the cluster analysis, but when looking at the full set of genes listed in the matching supplemental figures there are definitely some genes strongly changing, just not necessarily the same ones. This may be due to differences in the myogenic state of the material studied: while Bakay and colleagues used muscle biopsies containing terminally differentiated muscle fibres, we focused on the earlier stages of myogenesis by *in vitro* differentiating cultured myoblasts obtained from muscle biopsies. Despite this, it is important to note that while different genes may be affected, most of the same pathways were highlighted in both Bakay’s and our study. Collagens COL6A1, COL6A2, COL6A3, COL12A1 are linked to Bethlem Muscular Dystrophy ^42–45^ and, interestingly, all these collagens were upregulated in Group 1 patient cells and downregulated in Group 3 patient cells (**Figure S5**). Matrix metalloproteinases were also altered with 13 out of 28 matrix metalloproteinases exhibiting changes in at least one of the comparisons and all patients had multiple of these genes altered (**Figures 4** **and S5**). Notably the metalloproteinase MMP1 (collagenase I), which has been proposed to resolve fibrotic tissue ^46^, was downregulated in all but one patient, as well as in Bakay EDMD samples. Likewise, multiple genes associated with fibrosis from FibroAtlas (**Figures 4** **and S6**) and with inflammation that would support fibrosis such as cytokine (**Figures 4** **and S7**) and INF-gamma signaling (**Figures 4** **and S8**) were affected in all patients. In fact, out of 941 genes in FibroAtlas there were 542 altered between all the patients. Heatmaps of gene clusters with similar expression patterns are shown in Figure 4, but more detailed individual panels with all gene names listed are shown in **Figures S5-S11**. A few genes that stand out for their functions within the INF-gamma signaling pathway include IRF4 that is a regulator of exercise capacity through the PTG/glycogen pathway ^47^ and ILB1 that helps maintain muscle glucose homeostasis ^48^ such that both could also feed into the metabolic pathways altered.

**Figure 4.**
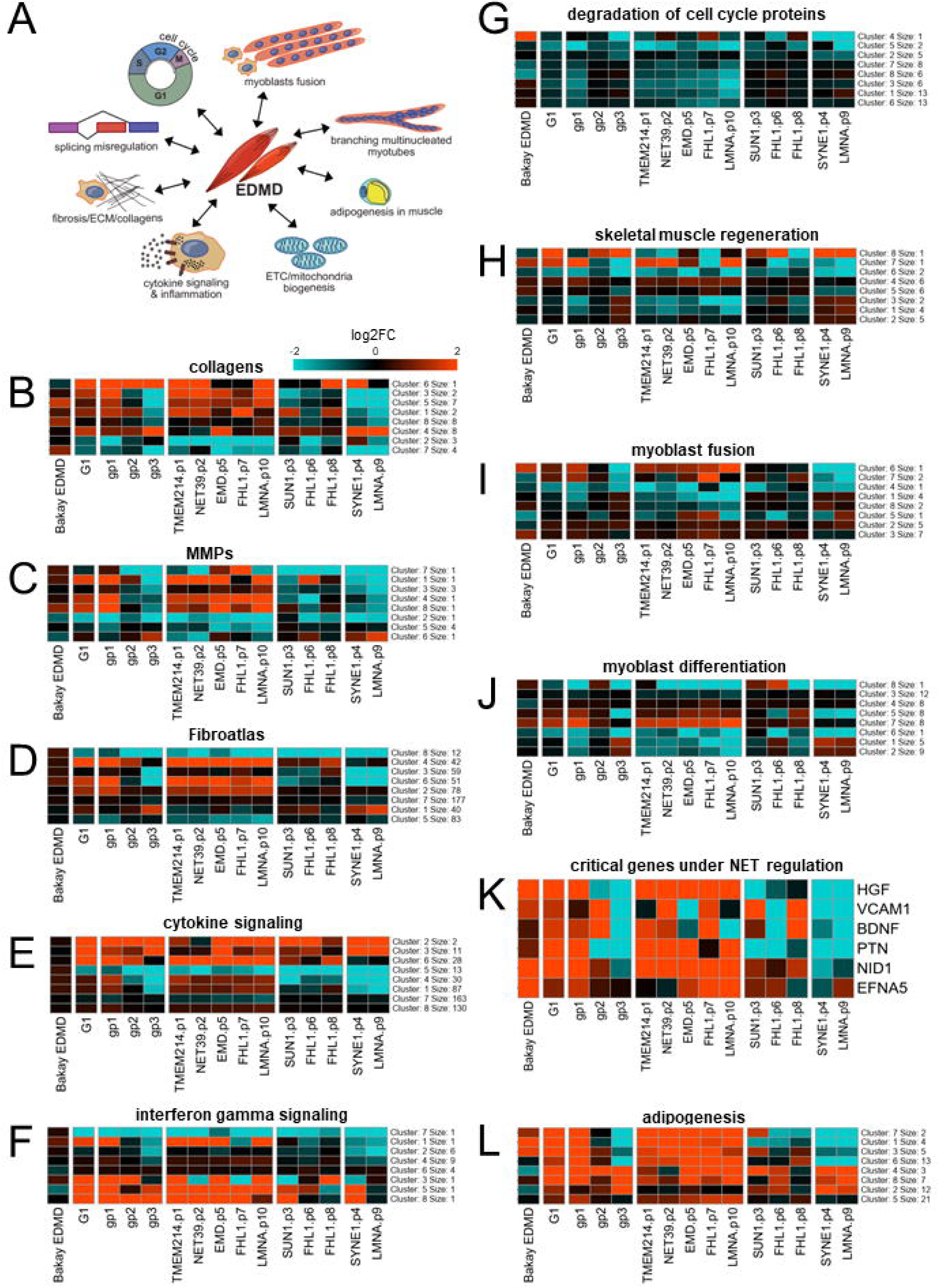
Genes changing expression within functional groups supporting ECM/ fibrosis/ myotube fusion. A. Summary of altered functions in EDMD. B. Heatmap showing gene expression changes for collagens. C. Heatmap showing gene expression changes for matrix metalloproteinases. D. Heatmap showing gene expression changes for general proteins involved in fibrosis according to FibroAtlas (http://biokb.ncpsb.org.cn/fibroatlas). E. Heatmap showing gene expression changes for cytokine signaling proteins (GO-BP, GO:0019221). F. Heatmap showing gene expression changes for interferon-gamma signaling proteins (WikiPathways, WP619). G. Heatmap showing downregulation of genes involved in the degradation of cell cycle proteins. GSEA identified downregulation in EDMD patients of REACTOME hsa-1741423 (APC-C mediated degradation of cell cycle proteins) and hsa-187577 (SCF/SKP2 mediated degradation of p27/p21) (Fig. 2C). Both genesets were combined. H. Heatmap showing gene expression changes for skeletal muscle regeneration proteins (GO-BP, GO:0043403). I. Heatmap showing gene expression changes for myoblast fusion (GO-BP, GO:0007520). J. Heatmap showing gene expression changes for myoblast differentiation (GO-BP, GO:0045445). K. Heatmap showing gene expression changes for specific genes functioning in myotube fusion that are under genome-organizing NET regulation. L. Heatmap showing the upregulation of many pro-adipogenic genes (Wikipathways, WP236). All heatmaps were generated from differentially expressed genes in at least one of Bakay’s EDMD, G1 analysis (all 10 patients analyzed as a single group), or G3 analysis (each of the three subgroups analyzed separately). K-means unsupervised hierarchical clustering was used to summarize the heatmap into 8 clusters of roughly coexpressing genes. Full heatmaps displaying gene names are provided in **Figures S5-S11**. The number of genes per cluster is indicated on the right of each heatmap. Red and blue indicate upregulation and downregulation respectively.

Another subpathway critical for myogenesis and the timing and integrity of myotube fusion is cell cycle regulation. Cell cycle defects could lead to spontaneous differentiation and were previously reported in myoblasts from EDMD patients and in tissue culture cell lines expressing emerin carrying EDMD mutations which could lead to depletion of the stem cell population ^41; 49^. All tested EDMD patients exhibited downregulation of multiple genes involved in the degradation of cell cycle proteins (**Figures 2**, **4**, and **S8**) which could indicate an uncoupling of the joint regulation of cell cycle and myogenesis program ^50^, for example cells starting to fuse when they should still be dividing or *vice versa*.

Other pathways in addition to ECM deposition directly associated with myogenic differentiation, myoblast fusion, and muscle regeneration were also altered in all patients (**Figures 4** **and S9-S10**), though, again, no single gene in these pathways was altered in the same way in all patients’ cells. Poor differentiation and myotubes with nuclear clustering were observed in differentiated EDMD myoblast cultures ^14^ and in the mouse C2C12 differentiation system when EDMD-linked NETs were knocked down ^17^.

Previous work using C2C12 cells identified six genes whose products are required in the early differentiation stages and were under the regulation of muscle-specific genome-organizing NETs ^17^. These genes (*NID1*, *VCAM1*, *PTN*, *HGF*, *EFNA5*, and *BDNF)* are critical for the timing and integrity of myotube fusion and need to be expressed early in myoblast differentiation but shut down later or they inhibit myogenesis ^51–55^. All six genes were misregulated in at least 5 but none were affected in all patients (**Figure 4**). In general terms, these genes were upregulated in Group 1, downregulated in Group 3, and mixed in Group 2. All six genes were upregulated in Bakay’s EDMD data, although only *NID1* and *HGF* were statistically significant at 5% FDR. Both were upregulated only in Group 1 and downregulated in Group 3. *PTN* showed a similar pattern of expression as *HGF* although the only statistically significant changes were for upregulation in Group 1.

Several myogenic signaling pathways were altered such as MAPK, PI3K, BMP and Notch signaling and several alternate differentiation pathways were de-repressed such as adipogenesis that could disrupt myotube formation and function (**Figure 2** **and Table S3**). Myogenesis and adipogenesis are two distinct differentiation routes from the same progenitor cells and whichever route is taken the other becomes repressed during normal differentiation ^56; 57^. We previously showed that knockout of fat-or muscle-specific genome organizing NETs yield de-repression of the alternate differentiation pathway ^17^;^58^ and the Collas lab showed that Lamin A lipodystrophy point mutations yield de-repression of muscle differentiation genes in adipocytes ^59^. We now find here that adipogenesis genes are upregulated in both Bakay’s EDMD data and our data (**Figure 2**). This is especially prominent for the five patients in Group 1 while Group 3 showing strong downregulation of a subset of the same genes and Group 2 broadly looking like an intermediate of the other two groups (**Figures 4** **and S11-S12**), and thus could also contribute to the metabolic defect differences between patients.

### Splicing Pathways Uniformly Altered in EDMD Patients Yield Loss of Muscle-Specific Splice Variants

Among the downregulated functional categories, mRNA splicing stood out with many genes uniformly downregulated in all patient samples (**Figures 2 and S13**). Because of that we decided to investigate various subcategories and we found that there was a striking and uniform upregulation of factors supporting alternative splicing while constitutive splicing factors involved in spliceosome assembly and cis splicing are downregulated (**Figures 5** **and S14**). Expression changes of as little as 10% (log2FC > 0.1) have been shown to result in biologically relevant changes for vital proteins like kinases and splicing factors ^60^. We thus assume that up- and down-regulation of whole spliceosome subcomplexes even in low log2FC ranges lead to significant splicing misregulation. Notably, snRNAs of the U1 spliceosomal subcomplex, responsible for 5’ SS recognition, constitute as much as 20% of all downregulated splicing factors (log2FC > |0.1|, RNU1s and RNVU1s). Interestingly, a similar sharp cut-off between alternative and constitutive splicing has been reported in myotonic dystrophy (DM1/DM2) with similar genes being affected, namely *CELF*s, *MBNL*s, *NOVA*, *SMN1/2* and *SF3A1*, among others ^61^. DM1 is one of the best studied splicing diseases and shares typical muscular dystrophy symptomology with EDMD. Moreover, a number of splicing changes in DM1 and DM2 also occur in other muscular dystrophies ^62^. Of note, *MBNL3* is 4-fold transcriptionally upregulated in EDMD compared to controls. Its protein product impairs muscle cell differentiation in healthy muscle and thus needs to be downregulated upon differentiation onset ^63^.

**Figure 5.**
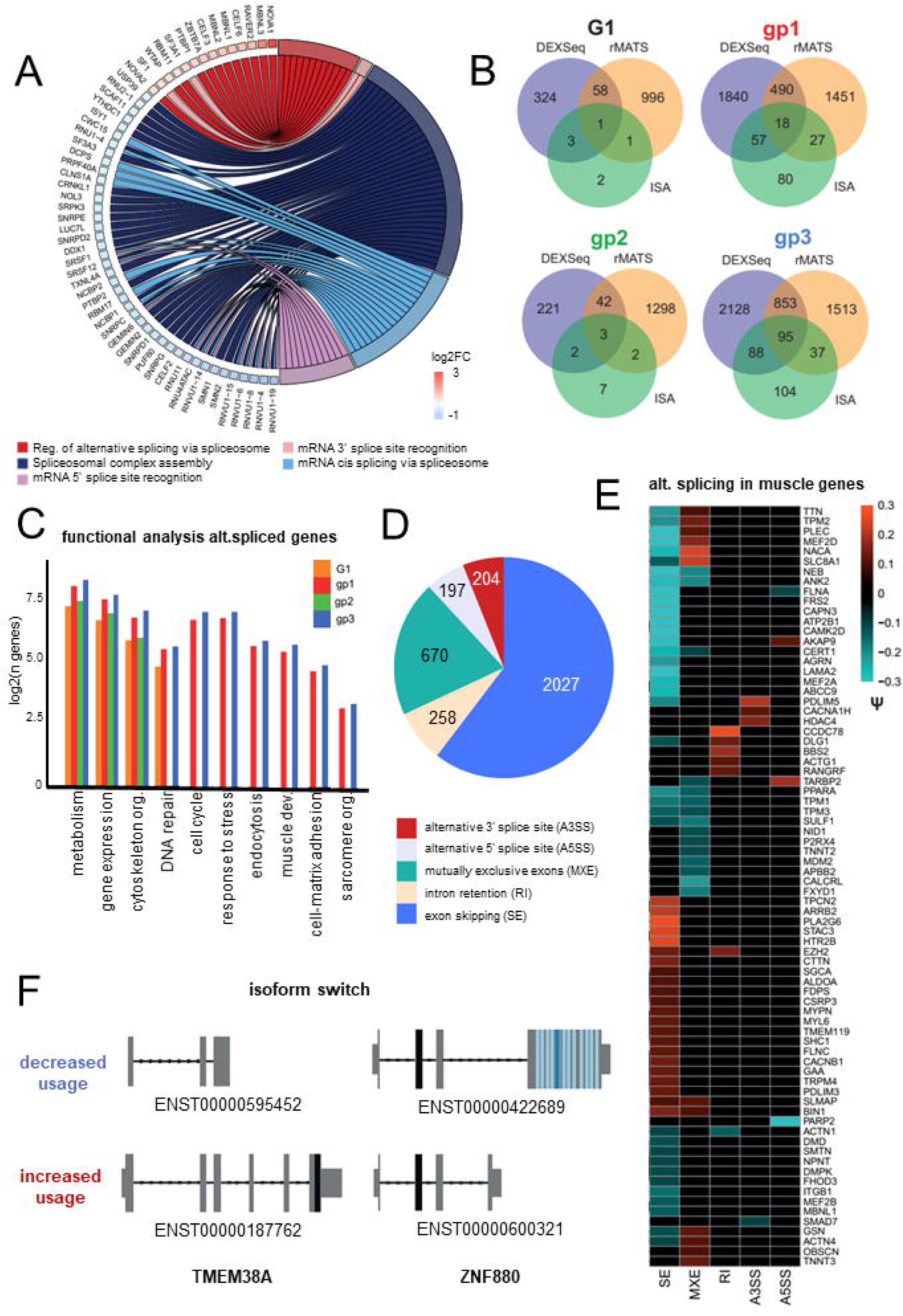
Analysis of splicing defects reveals loss of muscle-specific splice variants. A. GOchord plot for misregulated splicing factors indicates the primary change is upregulation of alternative splicing and downregulation of constitutive splicing. Plot for gp1 is shown as a representative. DE genes log2FC > |0.15| and p < 0.05. B. Venn diagrams of alternatively spliced genes predicted by rMATS, DEXSeq and ISA for all samples (G1) and the three subgroups (gp1, gp2 and gp3). Overlap between all three methods indicates genes with exon skipping events that lead to annotated isoform switches. C. Bar chart for functional pathways reveal an enrichment in altered splice variants for pathways associated with metabolism, gene expression, cytoskeleton organization, DNA repair, proliferation/ differentiation, stress, ECM/ fibrosis/ and sarcomere structure. Number of genes detected displayed as log2. E. Pie Chart of significant AS events with psi > |0.1| in gp1 detected with rMATS, which were used to scan for muscle specific splice variants, shown as Heatmap. AS events SE = exon skipping, MXE= mutually exclusive exons, RI = intron retention, A3SS = alternative 3’ splice site, A5SS = alternative 5’ splice site. F. Isoform switches as analyzed using isoformSwitchAnalyzer for ZNF880 and TMEM38A. ZNF880 shows the same isoform switch in all groups with preferential use of the shorter isoform, that contains the KRAB domain (black), but not the zinc finger domain (blue). TMEM38A shows a clear switch from the muscle isoform to a shorter isoform.

Next, we performed splicing analysis to determine whether mis-splicing could drive some of the pathway alterations observed in the EDMD samples. For this purpose, we used three different methods: DEXSeq analyses exon usage, rMATS provides information about the five most common alternative splicing (AS) events and isoform Switch Analyzer (ISA) indicates which splicing events lead to annotated isoform switches. This revealed varying amounts of alternatively spliced genes in all samples and the three subgroups (**Figure 5**). Since every method focuses on a different event type/ aspect of splicing, a higher amount of unique than overlapping genes is to be expected. Accordingly, an overlap of all three methods indicates genes with exon skipping events that lead to annotated isoform switches. The numbers of mis-spliced genes overlapping between the three algorithms was only 1 gene, *ZNF880*, in G1. In contrast, when analyzing Group 1, 2, and 3 separately, each patient grouping had many mis-spliced genes identified by all three algorithms with 18 mis-spliced genes in the intersect for Group 1, the group including half of the patients, and as much as 95 in Group 3 (**Table S4**). These genes include Nesprin 3 (*SYNE3*), the splicing factor kinase *CLK1* and the chromatin regulator *HMGN3*, all of which are potentially contributing to EDMD, given their functions. All results can be found in **Table S5**. The rMATS analysis includes five AS events: exon skipping (SE), intron retention (RI), mutually exclusive exons (MXE), alternative 3’ splice site (A3SS) and alternative 5’ splice site (A5SS) usage. Using this comprehensive dataset, we searched for AS events that are significantly differentially used (percent-spliced-in-(psi)-value > |0.1| and p-value < 0.05, **Table S5**). Comparing AS event inclusion between control and EDMD samples, we find thrice as much intron retention in EDMD, while all other events are similarly included as excluded. We hypothesize that this could be a result of downregulated U1 snRNAs which are necessary for proper spliceosome assembly. Supporting the likely importance of the splicing pathway to the EDMD pathomechanism, pathway analysis on these genes revealed a strong enrichment for pathways associated with metabolism, gene expression and the cytoskeleton (**Figure 5**). Moreover, Group 1 and Group 3 display an enrichment for myogenesis and muscle contraction. Using a custom-made set of genes either specific or relevant for muscle development and structure (see methods), we then scanned all significant and differential AS events (**Figures 5** **and S15**). Notably, mis-splicing led to the absence of many muscle-specific splice variants (**Figure 5**), among them vital muscle structural genes like *TTN*, *TNNT3*, *NEB*, *ACTA4* and *OBSCN* as well as developmental regulators of the MEF2 family. Importantly, many of them are linked to a variety of muscular dystrophies. For example, *TTN*, *CAPN3*, *PLEC*, and *SGCA* are linked to Limb-Girdle muscular dystrophy ^64–69^, *DMD* is linked to Duchenne Muscular Dystrophy and Becker muscular dystrophy ^70; 71^, and BIN1, TNNT2/3 and MBNL1 are mis-spliced in myotonic dystrophy ^72^.

Intriguingly, one of the mis-spliced genes which also displays an isoform switch in Group 3 is *TMEM38A* that has been linked to EDMD ^10^. The altered splicing map for *TMEM38A* reveals that not only is its expression highly elevated in EDMD patients (log2FC = 2.9), but also that the protein-coding isoform displays a higher usage relative to abundance compared to the non-coding isoform (**Figure 5**) Many other notable mis-spliced genes are involved in myotube fusion such as the previously mentioned *NID1* that is under spatial genome positioning control of NET39, another of the genome organizing NETs causative of EDMD. Most compellingly, three mis-spliced genes having to do with myogenesis/ myotube fusion had muscle-specific splice variants absent in all patients (G1 rMATS). These were *CLCC1* whose loss yields muscle myotonia ^73^, *HLA-A/B* that disappears during myogenesis and is linked as a risk factor for idiopathic inflammatory myopathies ^74; 75^, and *SMAD2* that shuts down myoblast fusion ^76^. The above examples were found in all patients within a particular Group, but not always amongst all patients from the study or determined by all algorithms; however, there were also some mis-spliced genes that are potentially even more interesting because they were mis-spliced in all patients and with all 3 algorithms yielding the same results. One of these was *ZNF880*. While overall transcript numbers remained similar, the isoform predominantly expressed in control cells, ENST00000422689, is strongly downregulated in Group 1 while the shorter isoform, ENST00000600321, is strongly upregulated (**Figure 5**). Interestingly, the dominant isoform in EDMD loses the zinc finger domain (light and dark blue) and is left with the repressive KRAB domain (black). Little is known about ZNF880 except that it has an unclear role in breast and rectal cancer ^77; 78^ and additional experiments are necessary to elucidate its role in EDMD.

### miRNA-Seq Analysis of EDMD Patient Cells

Changes in miRNA levels have been observed in a number of muscular dystrophies and are often used as biomarkers ^79; 80^, but a comprehensive investigation of miRNA levels in EDMD has thus far not been engaged. Thus, the *in vitro* differentiated EDMD patient cells used for the preceding analysis were also analyzed by miRNA-Seq. We identified 28 differentially expressed miRNAs with some variation among patients (**Figure 6**). We extracted their putative targets from the miRDB database (web resources) and selected those targets whose expression changed in the opposite direction of the miRNAs. Pathway analysis revealed misregulation of miRNAs largely associated with the same pathways that were misregulated from the RNA-Seq data e.g. metabolism, ECM/ fibrosis, and signaling/ differentiation (**Figure 6** **and Table S6**). More specifically for metabolism 9 of the misregulated miRNA were linked to metabolic functions and with only partial overlap another 9 linked to mitochondria function, for ECM/ fibrosis 19 of the misregulated miRNAs were linked to ECM and again with only partial overlap 10 to fibrosis and 13 to cytokines and inflammation. As noted before the ECM category in addition to potentially contributing to fibrosis is also relevant for myotube fusion along with cell cycle regulation that was targeted by 13 misregulated miRNAs and myogenesis that was targeted by 5 miRNAs. Several misregulated miRNAs had functions relating to alternative differentiation pathways with 4 relating to adipogenesis, 12 to neurogenesis, 17 to angiogenesis and for signaling there were 9 misregulated miRNAs affecting MAPK pathways, 8 for Akt signaling, 1 for JAK-STAT signaling, 5 for TGF-beta signaling, 2 for Notch signaling, and 3 for TLR signaling. Interestingly, some misregulated miRNAs were also reported as being linked to disease states such as miR-140-3p to dilated cardiomyopathy through its repressive effect on the integrin metalloproteinase gene *ADAM17* ^81^. As well as working within cells, miRNAs are often detected within a circulating exosomal microvesicle population that can be harvested from blood serum. This makes them especially attractive as potential biomarkers when compared to more invasive biopsies, but a much larger sample size together with more clinical informaiton will be required to clarify these as biomarkers.

**Figure 6.**
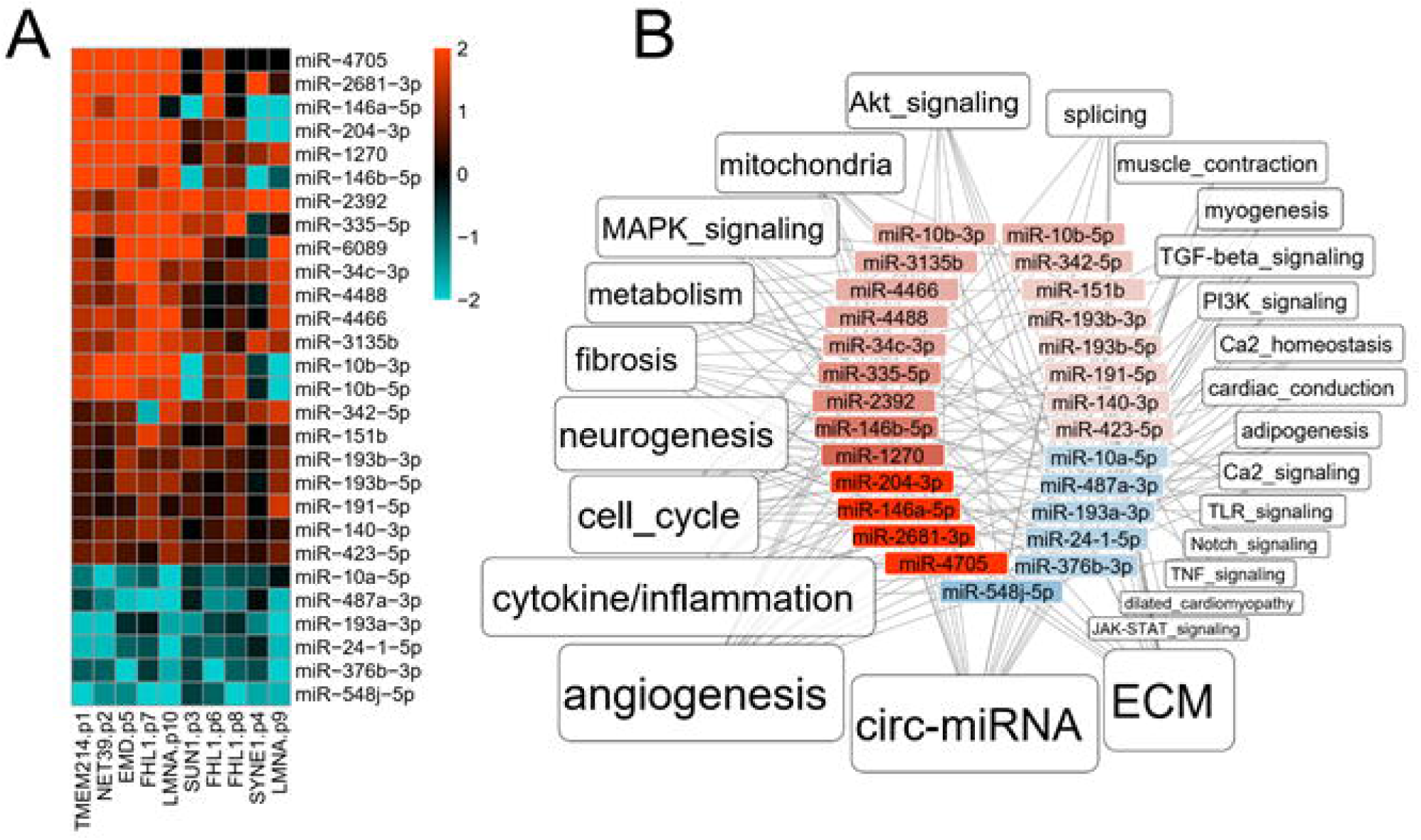
miRNA analysis. A. Heatmap of miRNAs that had altered levels in EDMD patient cells compared to controls. Red indicates upregulated and blue downregulated with intensity according to the log2FC values. B. Overview of functional categories linked to differentially expressed miRNAs in EDMD. The label size is proportional to the number of DE miRNAs associated with each category.

### EDMD Gene Expression Signature Suggests Relationships to Other Muscular Dystrophies

The earlier Bakay study analyzed patient samples from other muscular dystrophies for comparison to EDMD. We re-analyzed their data and used GSEA to determine how related the different muscular dystrophies are to our data. In order to visualize the results, for each disease we plotted the GSEA normalised enrichment score (NES) against the -log10(p-value). The higher on the y-axis the higher the confidence, while positive and negative correlations appearing to the right or to the left of the vertical, respectively. Thus, the further to the upper right the various Bakay disease sets are, the closer it is to the data derived from our set of 10 patients. When we looked at all patients as a single group, EDMD was the best match (**Figure 7**), with the highest score and lowest p-value (FDR 0.001). This indicates that despite the differences in the individual DE genes between the mature muscle data from Bakay’s EDMD geneset and our early *in vitro* differentiation geneset, a clear EDMD gene expression signature was displayed in our 10 patients. The next best match is Limb-Girdle muscular dystrophy 2A (LGMD2A), which is particularly interesting because *LMNA* mutations also cause Limb-Girdle muscular dystrophy 2B (LGMD2B) and this was even further distal in pathway analysis signatures from both our EDMD and the Bakay EDMD patients than Fascioscapulohumeral muscular dystrophy (FSHD) and Duchenne Muscular Dystrophy (DMD) patients. The differences in gene signatures that broke down the 10 patients into three EDMD patient subgroups could reflect an underlying cause of clinical disease spectrum or indicate that a group may not be adequately classified as EDMD. Therefore, we performed the same GSEA analysis on each subgroup separately. Group 1, which had both classic emerin and lamin A EDMD mutations showed an even better match with the Bakay EDMD group which was again very close to LGMD2A but also to DMD, Becker Muscular Dystrophy (BMD), FSHD and Limb-Girdle muscular dystrophy 2I (LGMD2I) (**Figure 7**). LGMD2B was still separate and closer to Juvenile Dermatomyositis (JDM). For Group 2, none of the diseases matched at FDR 5%, although the Bakay EDMD set remained the most like our set. Interestingly, two diseases exhibited an anti-correlation: DMD and BMD, which are both caused by mutations in the dystrophin gene *DMD*. In contrast, Group 3 appeared to be the most distinct and in many ways opposite to Group 1, which is a pattern that was often observed in the functional gene subsets analyzed (**Figures S5-S11**). Group 3 was anticorrelated with EDMD and most of the other muscular dystrophies, while the neurogenic Amyotrophic Lateral Sclerosis (ALS) appeared as the best match, possibly suggesting a neuronal bias in this group (**Figure 7**). This is further supported by the appearance of axonal neuropathy, ataxia, undergrowth, and speech problems in one of the two patients from this group (patient 4, *SYNE1*; **Table 1**), while none of the others exhibited any signs of neuropathy.

**Figure 7.**
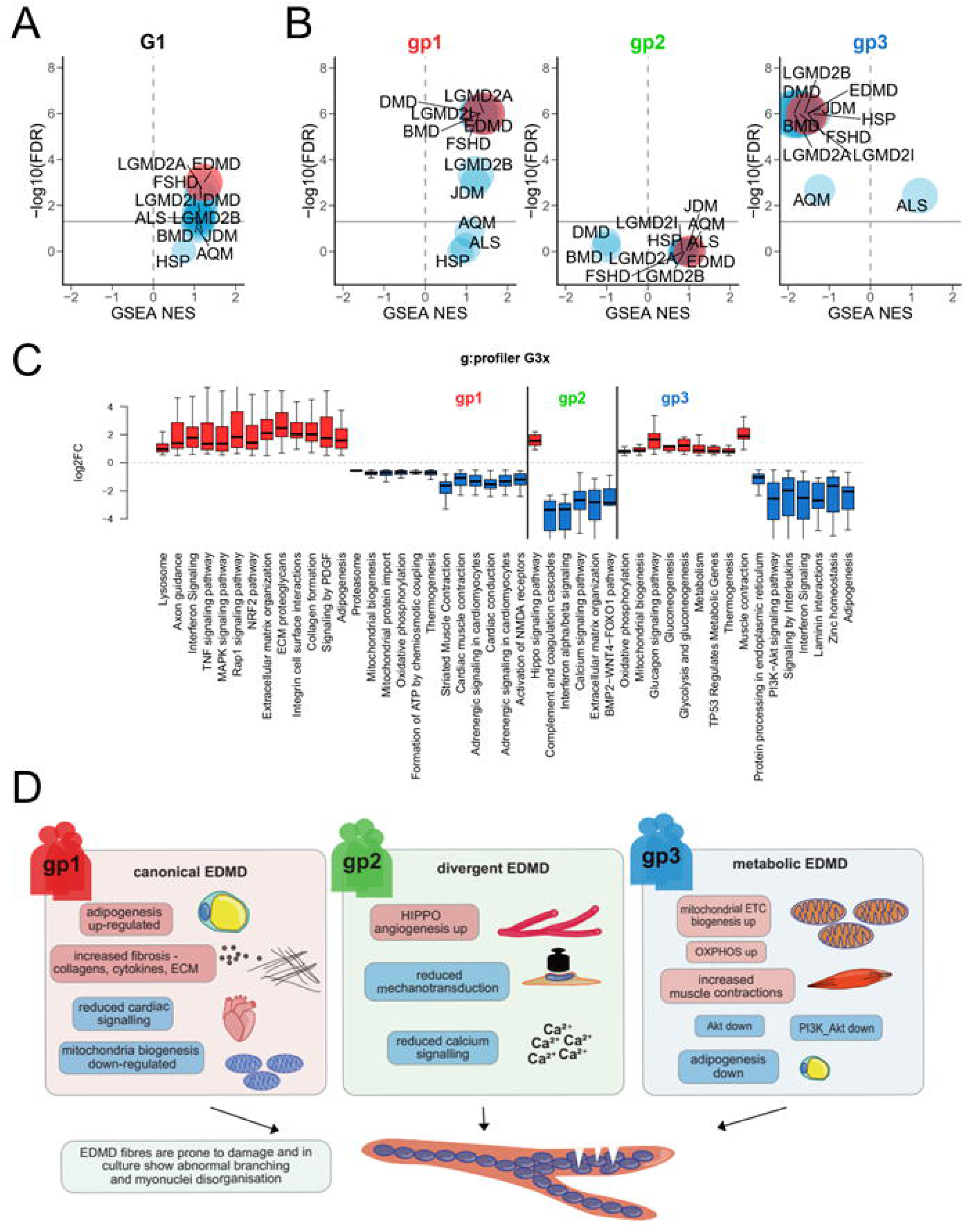
EDMD is distinct from other MDs. A. Scatterplots from GSEA analysis comparing Bakay data for several muscular dystrophies to the new patient data from this study (all 10 patients as a single group = G1). On the x axis, the Normalized Enriched Score (NES) is a measurement of the enrichment of the DE geneset identified in our study compared to each of the diseases in the Bakay study. The y axis shows the -log10(FDR), which is a measurement of statistical confidence. The grey horizontal line marks the 5% FDR threshold. Muscular dystrophies from the Bakay study are EDMD, Limb-Girdle muscular dystrophy 2A (LGMD2A), Limb-Girdle muscular dystrophy 2B (LGMD2B), Limb-Girdle muscular dystrophy 2I (LGMD2I), FascioScapuloHumeral muscular dystrophy (FSHD), Duchenne Muscular Dystrophy (DMD), Becker Muscular Dystrophy (BMD), Juvenile Dermatomyositis (JDM), Acute Quadriplegic Myopathy (AQM), Amyotrophic Lateral Sclerosis (ALS) and Hereditary Spastic Paraplegia (HSP). B. Same as A, for each individual patient subgroup. C. Box plot of log2FC values for differentially expressed genes within significantly enriched functional categories for each patient subgroup compared to the rest, using g:Profiler. D. Each patient subgroup was relatively more enriched for certain functional pathways than other subgroups, suggesting that treatments for example targeting different metabolic pathways for groups 1 and 3 might partially ameliorate some patient difficulties.

The relationship of the patient Groups segregated by gene signatures to potential differences in clinical presentation is underscored by the functional pathways enriched in each group over the others (**Figure 7**). Group 1 showed a strong enrichment of pathways associated with ECM and fibrosis, such as interferon signaling, TNF signaling, ECM organization, ECM proteoglycans, integrin cell surface interactions, collagen formation, and signaling by PDGF all upregulated. Adipogenesis was also particularly promoted in Group 1 compared to the others, and cardiac conduction defects were also highlighted. Group 2 was more uniquely associated with Hippo signaling and BMP2-WNT4-FOXO1 pathway and had fewer links to ECM and fibrosis. Group 3 was more uniquely associated with metabolism, particularly upregulation of oxidative phosphorylation, mitochondrial biogenesis, glucagon signaling pathway, gluconeogenesis, glycolysis and gluconeogenesis, metabolism, TP53 regulates metabolic genes, and thermogenesis pathways. This would suggest that Group 1 pathophysiology may have more characteristics of fibrosis and altered myofibers while Group 2 may have more differentiation or mechanosignaling defects and Group 3 more metabolic defects (**Figure 7**).

## Discussion

Attempts to identify the EDMD pathomechanism or clinical biomarkers purely through gene expression signatures are limited because there is too little uniformity in differential gene expression between all patients. We therefore engaged a functional pathway analysis using *in vitro* differentiated myotubes derived from 10 unrelated EDMD patients with known mutations in 7 EDMD-linked genes. While it is difficult to detect many individual genes that were uniformly changed in all patients, we found many pathways that were affected in all patients. Thus, although different genes may have been targeted in different patients, the same functional pathway would be disrupted and thus yield a pathology with similar clinical features. Many pathways were disrupted when we re-analyzed data from the previously published Bakay study and we postulated that, as they just analyzed mutations in two of the over two dozen genes linked to EDMD, analyzing a larger set of linked genes might narrow down the number of pathways to highlight those most relevant to EDMD pathophysiology. Indeed, when we considered a wider set of patients with mutations in 7 different genes the set of affected pathways narrowed to the point that we could identify four likely umbrella pathways.

These four umbrella pathways all make sense for contributing to or even driving the EDMD pathomechanism ^82^. Disruption of metabolism pathways from the gene expression analysis was consistent with the significantly reduced glycolysis and mitochondrial respiration output we showed in patient myoblasts compared to controls and it makes sense that this could lead to fatigue, weakness, and muscle atrophy. ECM changes and fibrosis pathways are consistent with pathology observed in EDMD and similarly could drive some of the initial pathology and, as fibrosis accumulates, contribute to disease progression. De-repression of genes from alternate differentiation pathways and defects in myogenesis through disrupted signaling pathways and cell cycle regulation could generate aberrant myotubes to yield pathology. Finally, the last disrupted pathway of splicing yields a loss of muscle-specific splice variants that could impact on all three preceding pathways.

There is much scope for intersection between the four highlighted pathways altered in all sampled EDMD patient cells. For example, amongst the de-repressed differentiation pathways was adipogenesis that could also impact on the metabolism pathway. Even amongst the few genes that were uniformly altered in all patients sampled, though not originally obvious, a more detailed reading of the literature leads to intersections with these pathways. For example, while the *MYH14* general upregulation did not make obvious sense for muscle defects since it is not part of the contractile machinery, it has been shown that a mutation in *MYH14* disrupts mitochondrial fission in peripheral neuropathy ^83^. Thus, *MYH14* could potentially feed into the mitochondrial deficits noted in the patient cells. Many of the miRNAs found to be altered in the patients feed into several of these pathways. For example, miR-2392 that is increased in all patients downregulates oxidative phosphorylation in mitochondria ^84^ but at the same time also is reported to promote inflammation ^85^. miR-140 that is up in all groups has roles in fibrosis through collagen regulation ^86^, is pro-adipogenic ^87^, and inhibits skeletal muscle glycolysis ^88^. miRNAs can also be used potentially prognostically between the different groups as for example miR-146a is upregulated in Group 1, unchanged in Group 2, and downregulated in Group 3. This miRNA has a strong effect on inflammation and has been implicated in fibrosis in the heart ^89^. Because there is so much functional overlap between miRNA targets and the pathways noted from the RNA-Seq analysis, it is unclear to what extent the gene expression changes observed could be indirect from the misregulated miRNAs. Nonetheless, there are 4 core functions targeted by multiple mechanisms that we argue are likely to be central to the core EDMD pathomechanism. Interestingly, the literature is filled with many examples of mutation or loss of different splicing factors causing muscle defects though no individual misspliced gene was identified as mediating these effects. Similarly, in myotonic dystrophy type 1 (DM1) there are many mis-spliced genes thought to contribute to the disease pathology ^72^. For example, the splicing factor SRSF1 that is down in most patients is important for neuromuscular junction formation in mice ^90^.

How so many genes become misregulated has not been experimentally proven, but for lamin, emerin, Sun1, nesprin, TMEM214 and PLPP7/NET39, the fact that mutations to all individually yield many hundreds of gene expression changes with considerable overlap strongly suggests that they function in a complex at the nuclear envelope to direct genome organization. It has already been shown that knockdown of Tmem214 and NET39 as well as several other muscle-specific NETs each alters the position and expression of hundreds of genes ^17^. Separately it was found that lamin B1 and the NET LAP2beta function together with two other proteins in a complex involved in tethering genes to the nuclear envelope ^91^ in fibroblasts and that emerin and lamins similarly function together with other proteins to tether genes in muscle cells ^92^. Thus, disruption of emerin, lamin A or any other component of these tethering complexes could yield sufficiently similar gene/ pathway expression changes to yield the core characteristic clinical features of EDMD. We propose that the different muscle-specific NETs give specificity to such a complex containing lamin A and emerin and that Sun1 and nesprin proteins can impact on these complexes through mediating mechanosignal transduction and FHL1 in interpreting such signals. Since 15% of all genes changing here were affected by at least one of the muscle-specific genome-organizing NETs that were tested by knockdown, this would provide a core set of genome organization and expression changes to cause the core EDMD pathology. Since the majority of genes affected by each NET tested was unique to that NET with the exception of Tmem214, this could account for other gene expression changes that drive the segregation into subgroups which could contribute to clinical variation. This interpretation is consistent with the numbers of genes changing for mutations in different nuclear envelope proteins (**Figure 1**). The patient cells with mutations in genome organizing NETs had much fewer genes changing than the patient cells with lamin A mutations. This could be because the lamin A mutations could disrupt multiple genome tethering complexes for different genome organizing NETs and thus alter expression of more genes. Alternatively, different lamin A mutations could preferentially yield disruption of complexes with particular genome-organizing NETs and thus patients with the same complex disrupted might share gene expression signature changes while those with different complexes disrupted might have less overlap so that this could account for different lamin A mutations segregating into different gene expression subgroups. In either case, the extreme differences in lamin A mutations gene expression profiles is not entirely surprising as different lamin mutations also exhibited large differences in studies of nuclear mechanics ^93^; so this could also impact on mechanosignal transduction. The Sun1 mutation may have affected fewer genes because its function in mechanosignal transduction is redundant with Sun2 while the Nesprin 1 mutation had more genes changing because it is more central to both mechanosignal transduction and to cell and nuclear mechanical stability.

The FHL1 mutations add another level of complexity to EDMD as there are several splice variants of FHL1 and only the B variant (ENST00000394155) targets to the nuclear envelope ^94^. That EDMD is a nuclear envelope disorder is underscored by the fact that none of the FHL1 mutations occur in exons found in the much shorter C variant (ENST00000618438) and the patient 8 mutation p.V280M is in an exon unique to FHL1B. Thus, the nuclear envelope splice variant is the only one that could yield pathology in all patients, though some of the variation could come from one of the patients also expressing the mutant A splice variant (ENST00000543669).

## Conclusions

While further work is needed to validate the correlations between the gene expression profile subgroupings and their clinical presentation and disease progression, our finding of such distinct gene expression signatures amongst clinically diagnosed EDMD patients argues that the currently used clinical phenotype spectrum umbrella of the EDMD classification may be too broad and it might be reclassified in more precise subtypes. What is clear is that the original classifications of EDMD subtypes based just on the mutated gene often allows for cases with very dissimilar gene signatures to be classified together while similar gene signature cases are classified as separate classes. The two lamin A mutations yielded changes in gene expression signatures that were far more different from one another than the group 1 lamin A mutation gene signature was from the TMEM214, NET39, emerin, and FHL1 mutation gene signatures. Similarly, the three FHL1 mutations yielded greater differences between them than these many genes in Groups 1 and 2. Thus, EDMD might be better classified by similarities in gene expression signatures than by the particular gene mutated. Regardless, these subtypes or separate disorders have distinctive gene expression signatures and miRNA signatures that could be used as biomarkers both diagnostically and perhaps prognostically. To get to that point will require a more comprehensive modern description of clinical Gestalt phenotypes including *e.g.:* imaging datasets and disease progression timelines deciphering unique groups. Importantly, the different pathways we found enriched for in each subgroup could be converted to clinical recommendations based on the much more conserved individual gene expression changes for each subgroup (**Figure 7**). For example, EDMD patients have been considered by some clinicians to be at risk for malignant hyperthermia ^95^, though a consensus was never achieved. Our data show that the 3 genes currently associated with malignant hyperthermia (*RYR1*, *CACNA1S*, and *STAC3*) are all misregulated in Groups 1 and 3, but not Group 2 (**Figure S16**). Thus, checking expression of these genes might indicate whether a patient is likely to be at risk or not. Finally, an additional new aspect coming up from our datasets is that it might be worth further investigating the role of splicing in muscle differentiation because it might be of wider relevance to muscular dystrophy beyond DM and EDMD.

## Data availability

Bakay *et al.* muscular dystrophy dataset is available at NCBI GEO with accession GSE3307. RNA-Seq and miRNA-Seq datasets will be deposited at NCBI GEO and made publicly available prior to publication.

## Acknowledgements

This work was funded by Muscular Dystrophy UK 18GRO-PG24-0248 and Medical Research Council MR/R018073 to ECS and Deutsche Forschungsgemeinschaft (DFG, German Research Foundation) – Projektnummer 470092532 to PM.

## Web resources

g:Profiler, https://biit.cs.ut.ee/gprofiler/gost

TissueEnrich, https://tissueenrich.gdcb.iastate.edu/

miRDB, http://mirdb.org

’smrnaseq’ miRNA-Seq analysis pipeline, https://nf-co.re/smrnaseq/1.1.0

miRTop, https://github.com/miRTop

NCBI GEO, https://www.ncbi.nlm.nih.gov/geo/

## Author Contributions

JIH processed patient samples for RNA- and miRNA-Seq and analyzed the data with assistance from SW. VT performed splicing analysis. LK-B, SH, and PM performed various metabolic analyses. RC helped with generation of figures and critical discussion. BS provided patient samples and inspiration. ECS designed the study and wrote the manuscript.

## Notes

### Competing Interest Statement

The authors have declared no competing interest.

