## Supplemental Figures S1-S16 and Supplemental legends for "Metabolic, Fibrotic, and Splicing Pathways Are All Altered in Emery-Dreifuss Muscular Dystrophy Spectrum Patients to Differing Degrees"

Figure S1

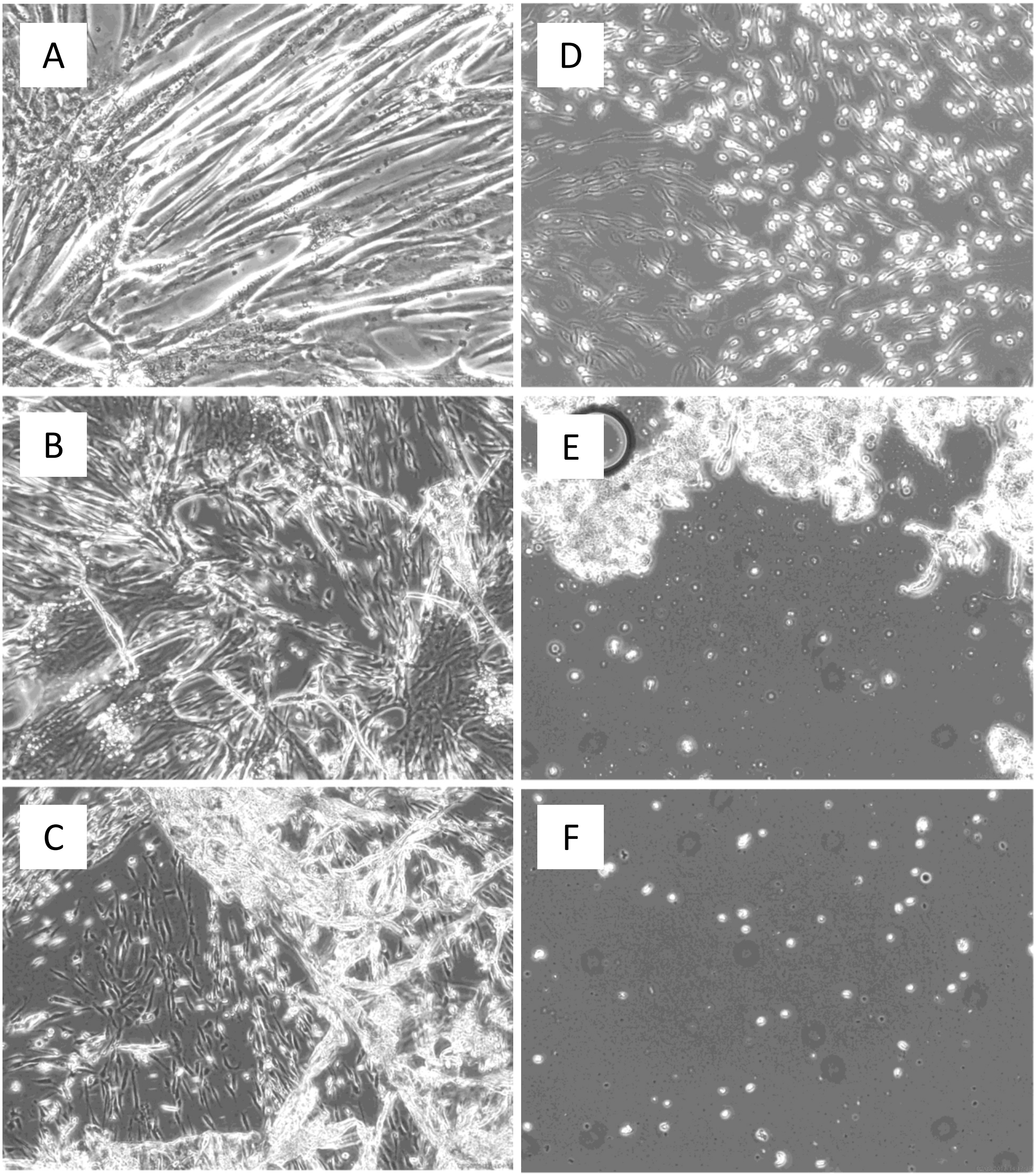

Figure S2

A

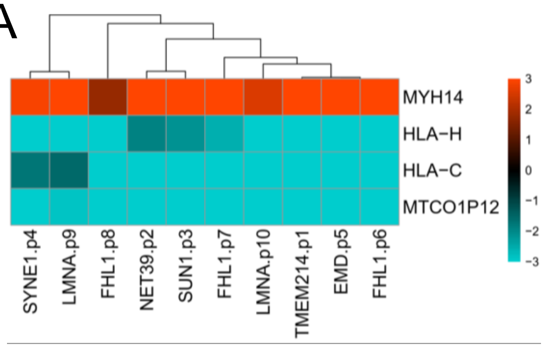


B

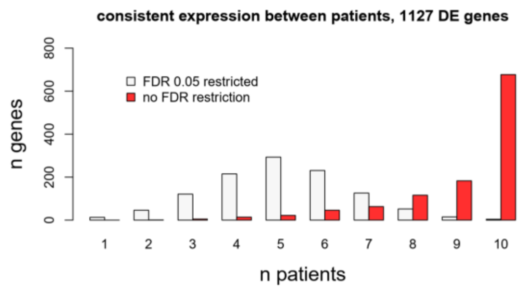


C

| FDR restriction | n patients | | | | | | | | | |
|---|---|---|---|---|---|---|---|---|---|---|
| | **1** | **2** | **3** | **4** | **5** | **6** | **7** | **8** | **9** | **10** |
| **0.05** | 13 | 46 | 121 | 215 | 293 | 231 | 126 | 52 | 15 | 4 |
| **1** | 0 | 1 | 5 | 14 | 22 | 46 | 63 | 116 | 183 | 677 |

D

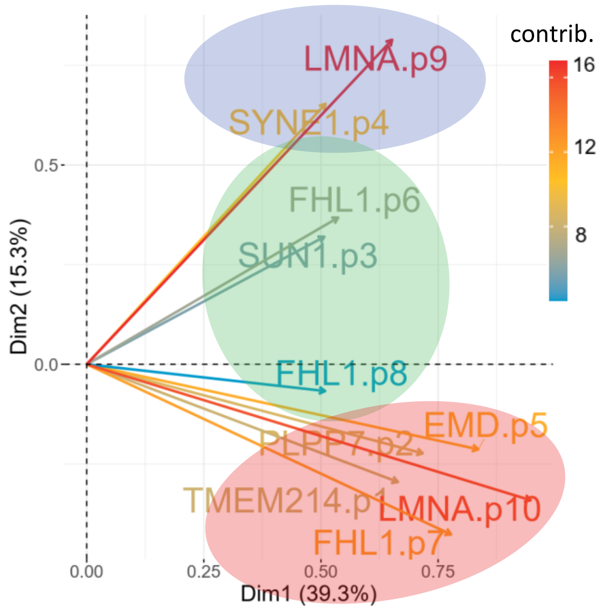

Figure S3

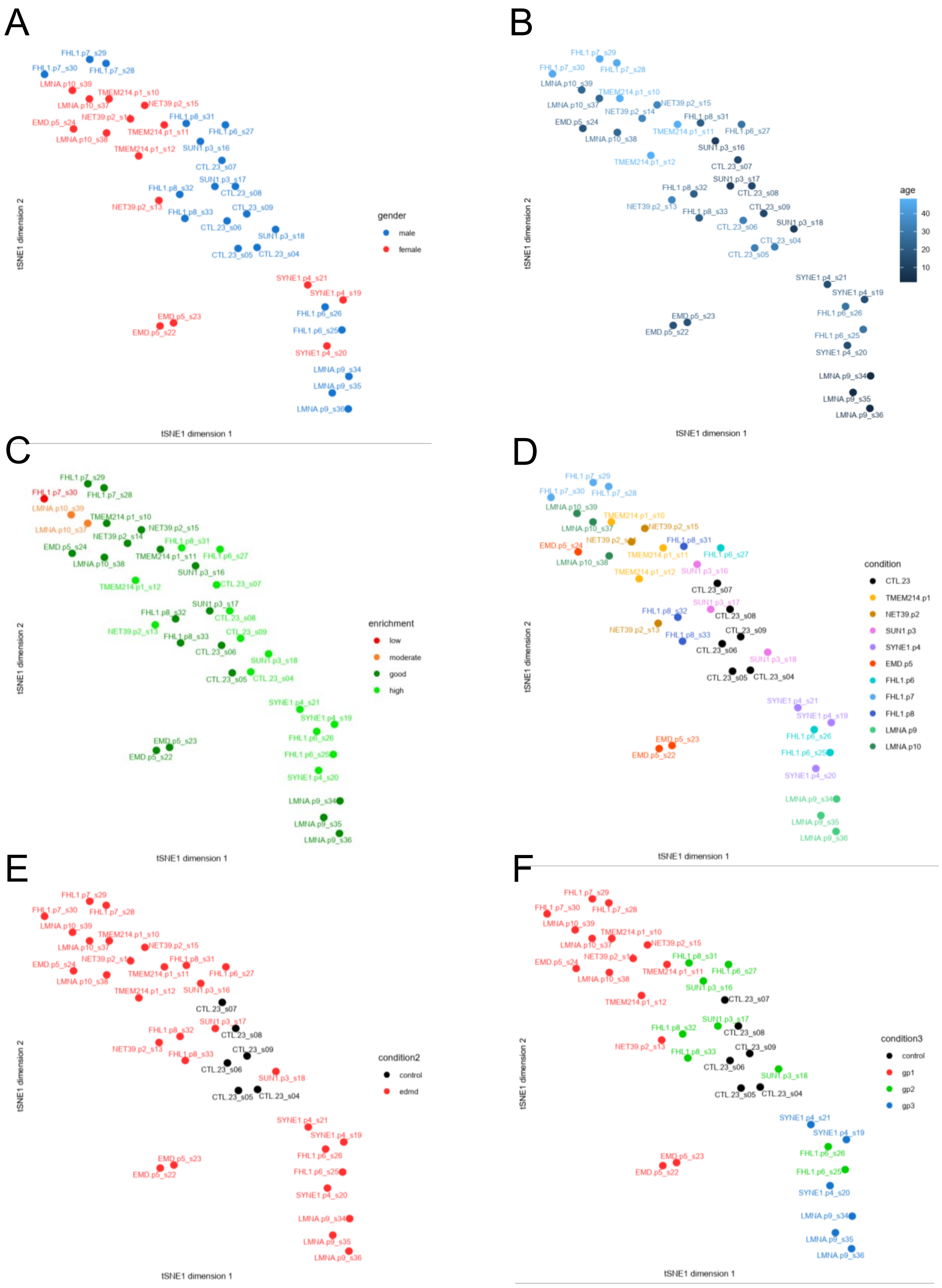
A
B
C
D
E
F
tSNE1 dimension 1
tSNE1 dimension 2
gender
male
female
age
40
30
20
10
enrichment
low
moderate
good
high
condition
CTL.23
TMEM214.p1
NET39.p2
SUN1.p3
SYNE1.p4
EMD.p5
FHL1.p6
FHL1.p7
FHL1.p8
LMNA.p9
LMNA.p10
condition2
control
edmd
condition3
control
gp1
gp2
gp3

Figure S4

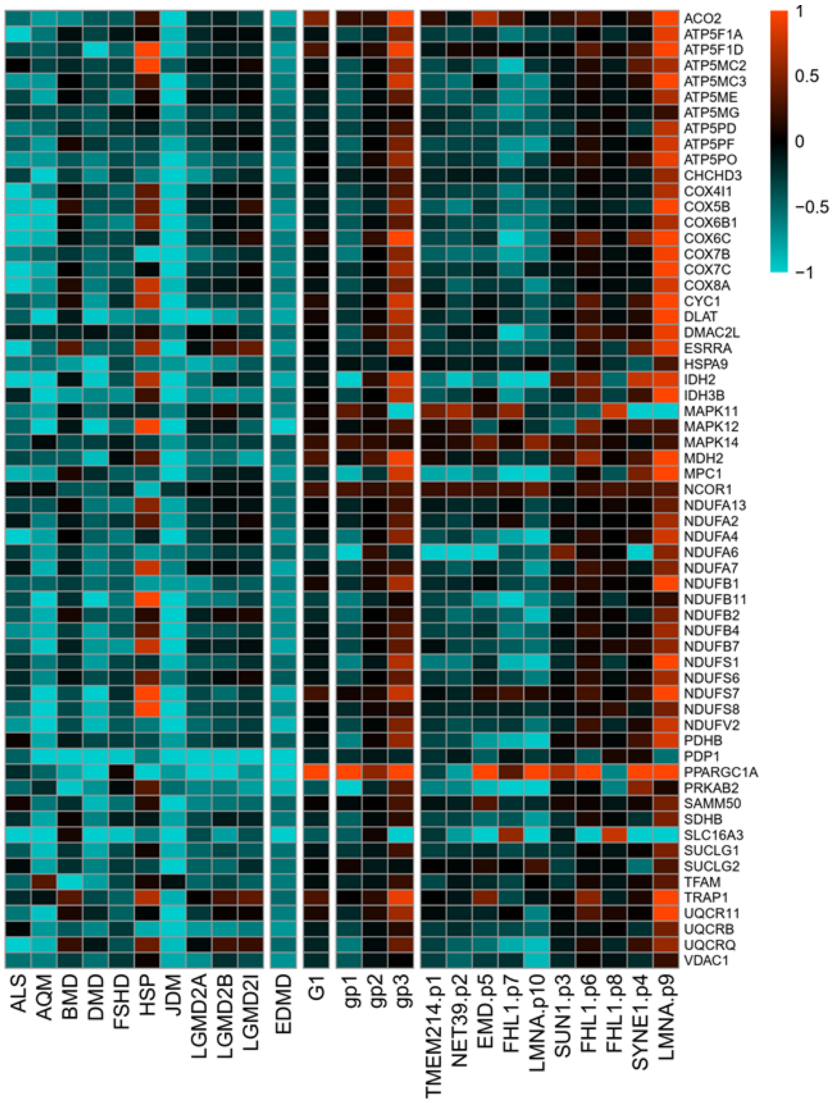
ACO2
ATP5F1A
ATP5F1D
ATP5MC2
ATP5MC3
ATP5ME
ATP5MG
ATP5PD
ATP5PF
ATP5PO
CHCHD3
COX4I1
COX5B
COX6B1
COX6C
COX7B
COX7C
COX8A
CYC1
DLAT
DMAC2L
ESRRA
HSPA9
IDH2
IDH3B
MAPK11
MAPK12
MAPK14
MDH2
MPC1
NCOR1
NDUFA13
NDUFA2
NDUFA4
NDUFA6
NDUFA7
NDUFB1
NDUFB11
NDUFB2
NDUFB4
NDUFB7
NDUFS1
NDUFS6
NDUFS7
NDUFS8
NDUFV2
PDHB
PDP1
PPARGC1A
PRKAB2
SAMM50
SDHB
SLC16A3
SUCLG1
SUCLG2
TFAM
TRAP1
UQCR11
UQCRB
UQCRQ
VDAC1
1
0.5
0
−0.5
−1
ALS
AQM
BMD
DMD
FSHD
HSP
JDM
LGMD2A
LGMD2B
LGMD2I
EDMD
G1
gp1
gp2
gp3
TMEM214.p1
NET39.p2
EMD.p5
FHL1.p7
LMNA.p10
SUN1.p3
FHL1.p6
FHL1.p8
SYNE1.p4
LMNA.p9

Figure S5

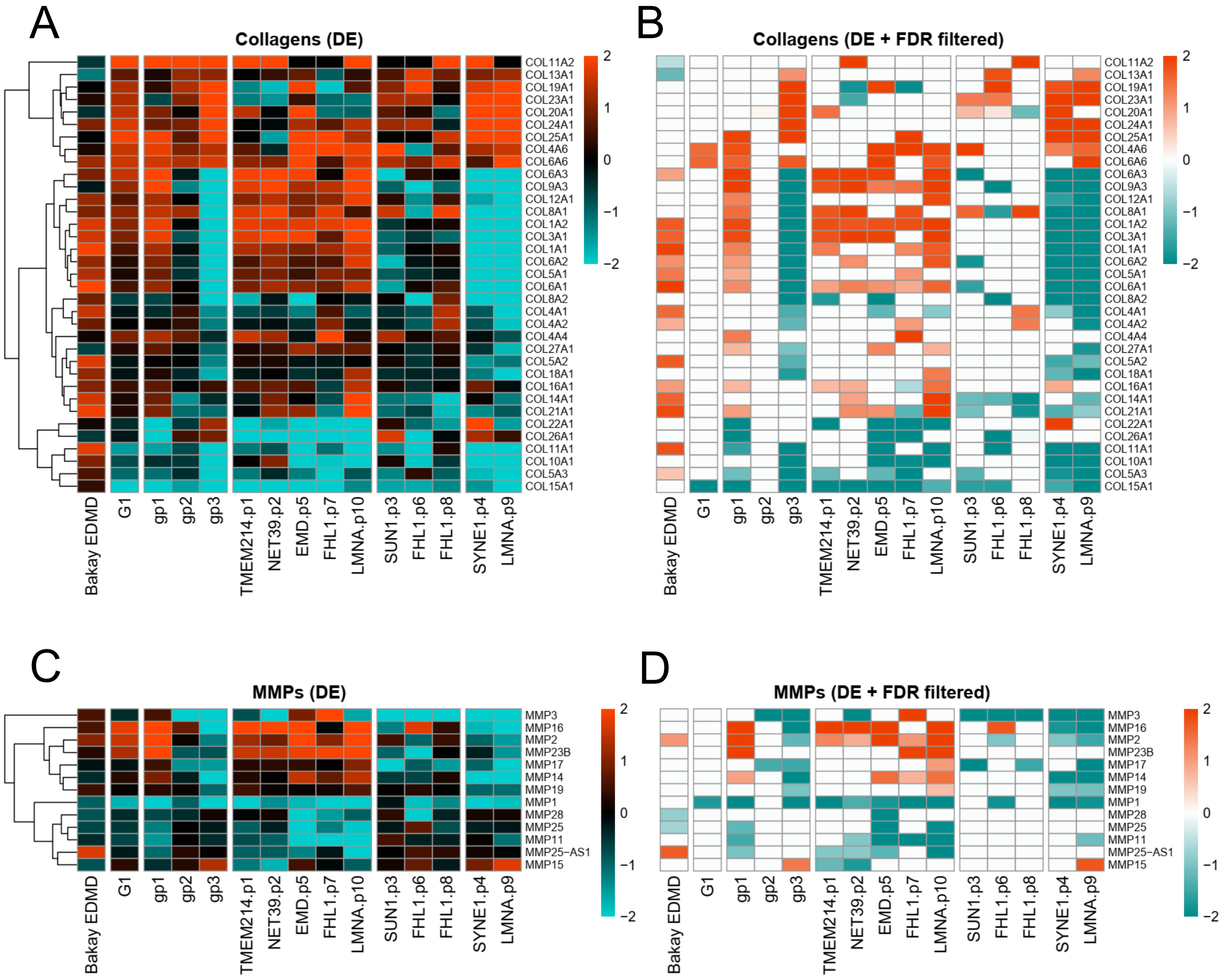
A
Collagens (DE)
B
Collagens (DE + FDR filtered)
COL11A2
COL13A1
COL19A1
COL23A1
COL20A1
COL24A1
COL25A1
COL4A6
COL6A6
COL6A3
COL9A3
COL12A1
COL8A1
COL1A2
COL3A1
COL1A1
COL6A2
COL5A1
COL6A1
COL8A2
COL4A1
COL4A2
COL4A4
COL27A1
COL5A2
COL18A1
COL16A1
COL14A1
COL21A1
COL22A1
COL26A1
COL11A1
COL10A1
COL5A3
COL15A1
Bakay EDMD
G1
gp1
gp2
gp3
TMEM214.p1
NET39.p2
EMD.p5
FHL1.p7
LMNA.p10
SUN1.p3
FHL1.p6
FHL1.p8
SYNE1.p4
LMNA.p9
C
MMPs (DE)
D
MMPs (DE + FDR filtered)
MMP3
MMP16
MMP2
MMP23B
MMP17
MMP14
MMP19
MMP1
MMP28
MMP25
MMP11
MMP25−AS1
MMP15
2
1
0
−1
−2

Figure S6

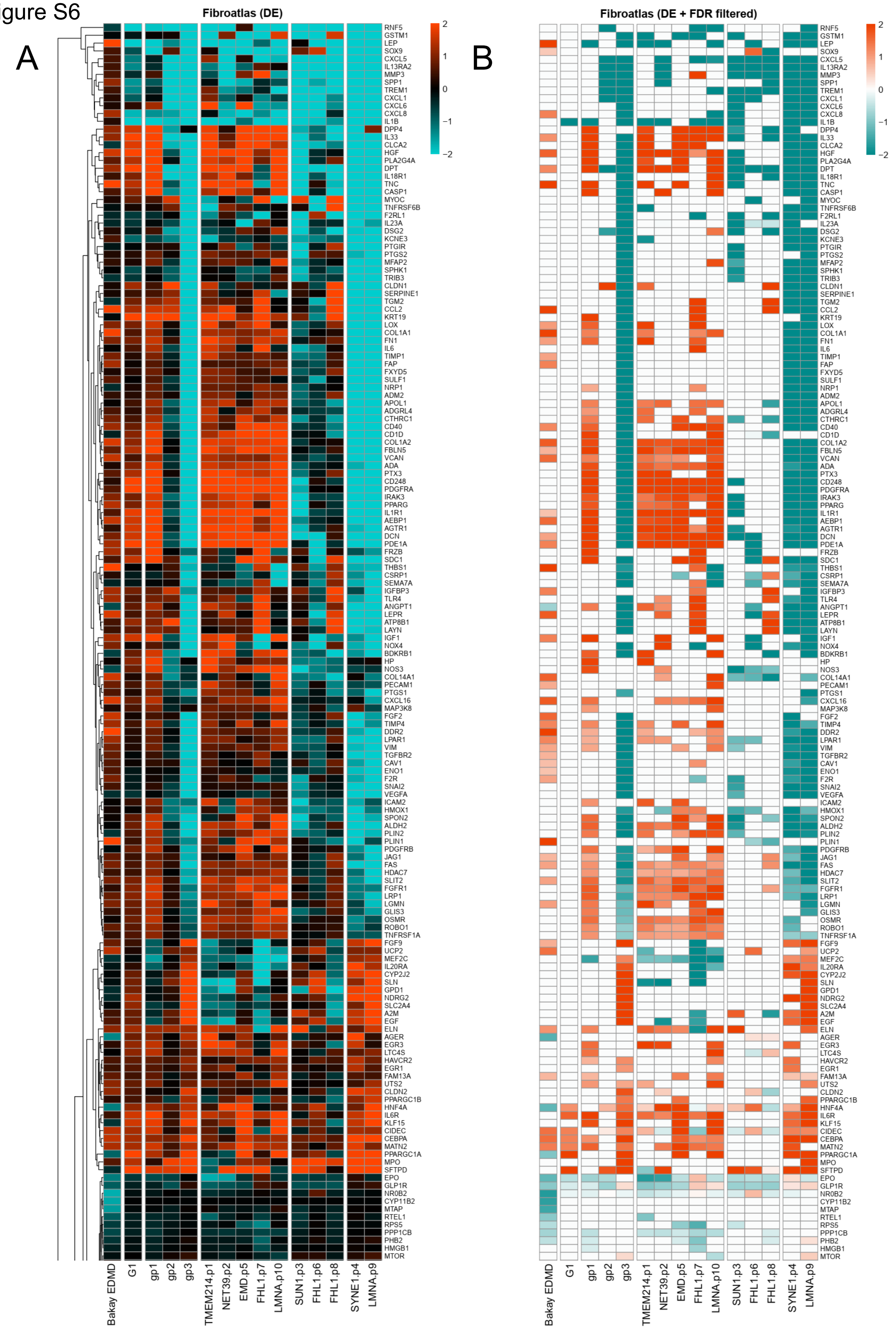
A
Fibroatlas (DE)
B
Fibroatlas (DE + FDR filtered)
2
1
0
−1
−2
RNF5
GSTM1
LEP
SOX9
CXCL5
IL13RA2
MMP3
SPP1
TREM1
CXCL1
CXCL6
CXCL8
IL1B
DPP4
IL33
CLCA2
HGF
PLA2G4A
DPT
IL18R1
TNC
CASP1
MYOC
TNFRSF6B
F2RL1
IL23A
DSG2
KCNE3
PTGIR
PTGS2
MFAP2
SPHK1
TRIB3
CLDN1
SERPINE1
TGM2
CCL2
KRT19
LOX
COL1A1
FN1
IL6
TIMP1
FAP
FXYD5
SULF1
NRP1
ADM2
APOL1
ADGRL4
CTHRC1
CD40
CD1D
COL1A2
FBLN5
VCAN
ADA
PTX3
CD248
PDGFRA
IRAK3
PPARG
IL1R1
AEBP1
AGTR1
DCN
PDE1A
FRZB
SDC1
THBS1
CSRP1
SEMA7A
IGFBP3
TLR4
ANGPT1
LEPR
ATP8B1
LAYN
IGF1
NOX4
BDKRB1
HP
NOS3
COL14A1
PECAM1
PTGS1
CXCL16
MAP3K8
FGF2
TIMP4
DDR2
LPAR1
VIM
TGFBR2
CAV1
ENO1
F2R
SNAI2
VEGFA
ICAM2
HMOX1
SPON2
ALDH2
PLIN2
PLIN1
PDGFRB
JAG1
FAS
HDAC7
SLIT2
FGFR1
LRP1
LGMN
GLIS3
OSMR
ROBO1
TNFRSF1A
FGF9
UCP2
MEF2C
IL20RA
CYP2J2
SLN
GPD1
NDRG2
SLC2A4
A2M
EGF
ELN
AGER
EGR3
LTC4S
HAVCR2
EGR1
FAM13A
UTS2
CLDN2
PPARGC1B
HNF4A
IL6R
KLF15
CIDEC
CEBPA
MATN2
PPARGC1A
MPO
SFTPD
EPO
GLP1R
NR0B2
CYP11B2
MTAP
RTEL1
RPS5
PPP1CB
PHB2
HMGB1
MTOR
Bakay EDMD
G1
gp1
gp2
gp3
TMEM214.p1
NET39.p2
EMD.p5
FHL1.p7
LMNA.p10
SUN1.p3
FHL1.p6
FHL1.p8
SYNE1.p4
LMNA.p9

Figure S6 (continued)

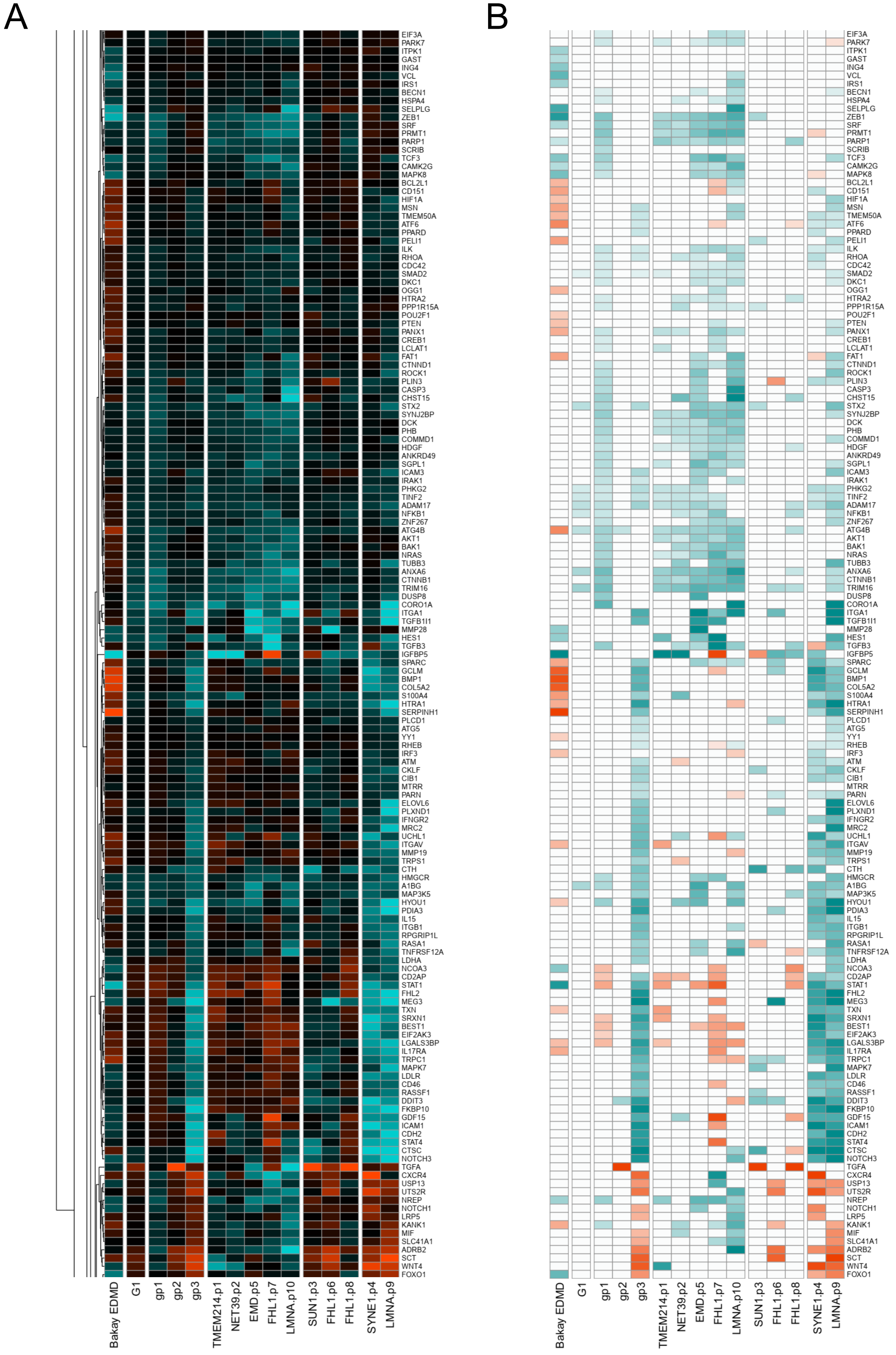

Figure S6 (continued)

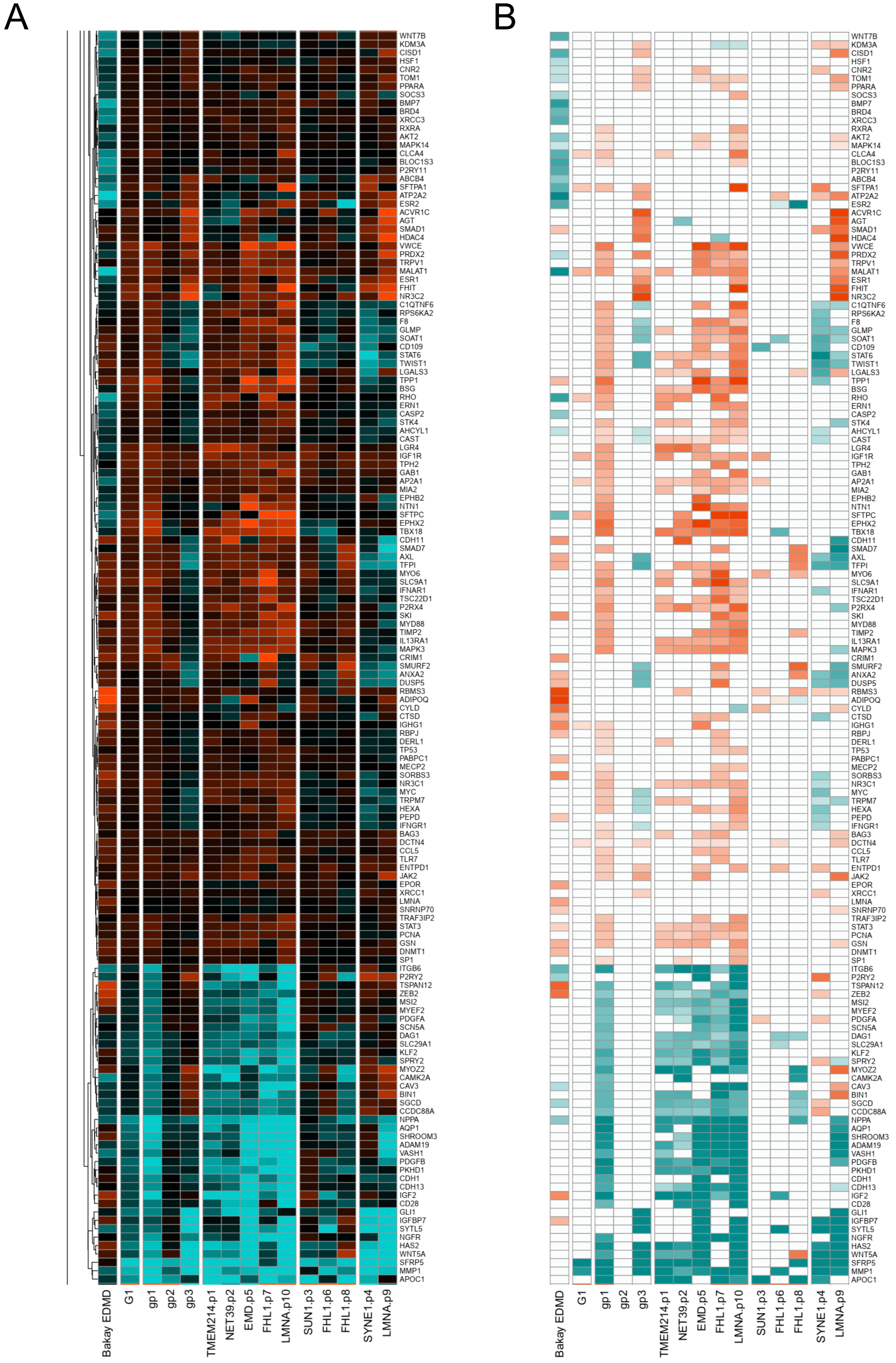

Figure S6 (continued)

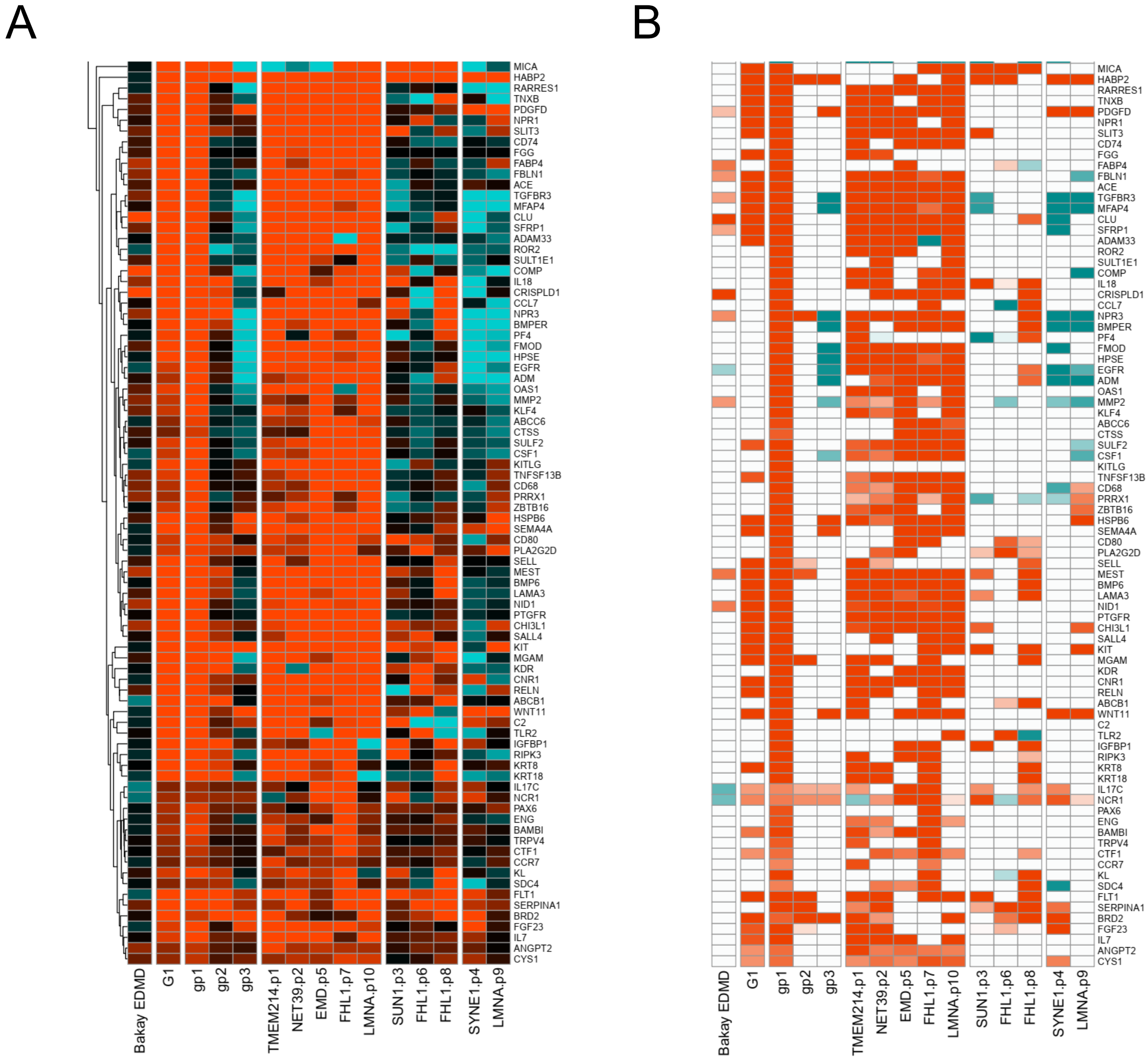

Figure S7

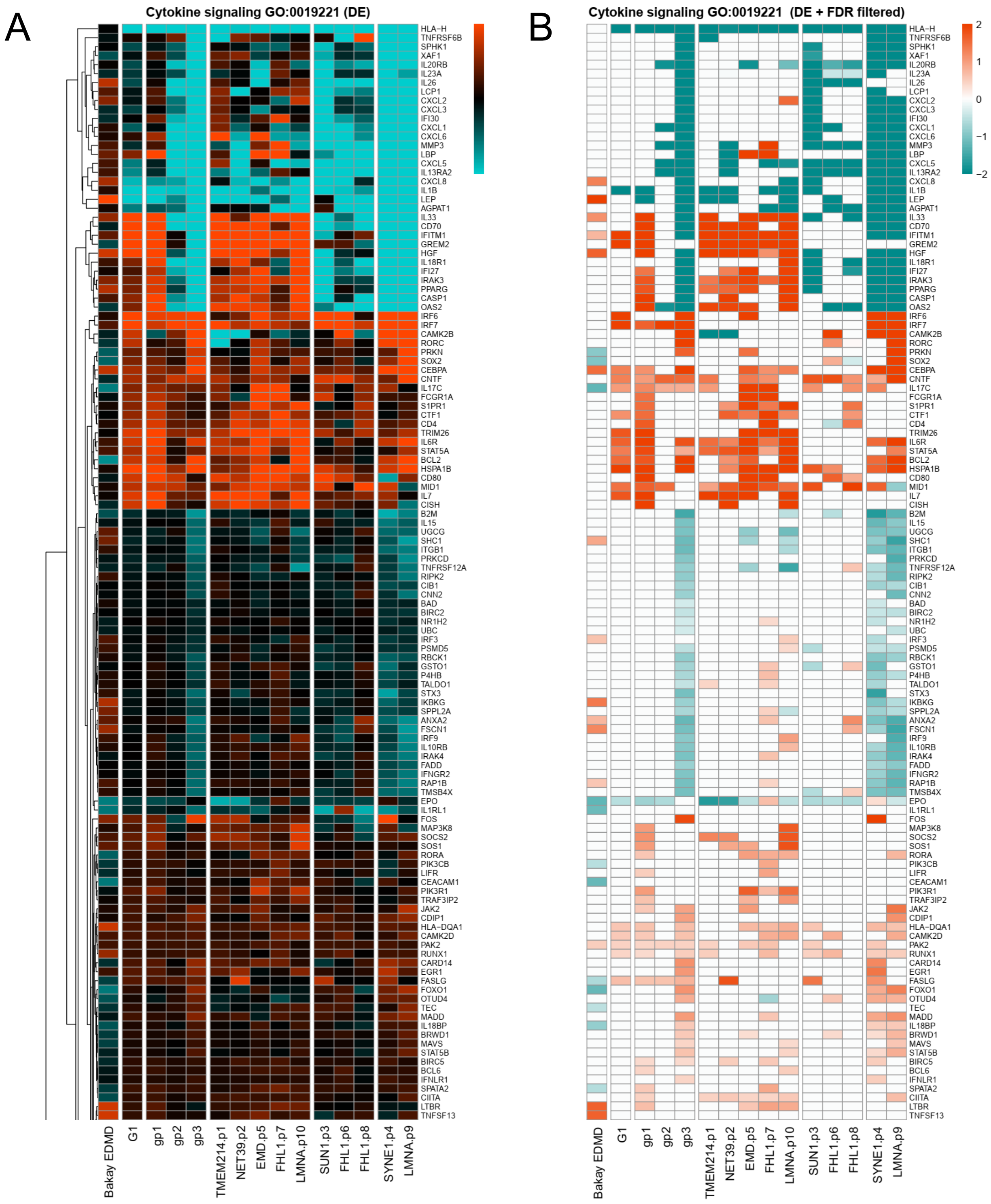
A
Cytokine signaling GO:0019221 (DE)
HLA−H
TNFRSF6B
SPHK1
XAF1
IL20RB
IL23A
IL26
LCP1
CXCL2
CXCL3
IFI30
CXCL1
CXCL6
MMP3
LBP
CXCL5
IL13RA2
CXCL8
IL1B
LEP
AGPAT1
IL33
CD70
IFITM1
GREM2
HGF
IL18R1
IFI27
IRAK3
PPARG
CASP1
OAS2
IRF6
IRF7
CAMK2B
RORC
PRKN
SOX2
CEBPA
CNTF
IL17C
FCGR1A
S1PR1
CTF1
CD4
TRIM26
IL6R
STAT5A
BCL2
HSPA1B
CD80
MID1
IL7
CISH
B2M
IL15
UGCG
SHC1
ITGB1
PRKCD
TNFRSF12A
RIPK2
CIB1
CNN2
BAD
BIRC2
NR1H2
UBC
IRF3
PSMD5
RBCK1
GSTO1
P4HB
TALDO1
STX3
IKBKG
SPPL2A
ANXA2
FSCN1
IRF9
IL10RB
IRAK4
FADD
IFNGR2
RAP1B
TMSB4X
EPO
IL1RL1
FOS
MAP3K8
SOCS2
SOS1
RORA
PIK3CB
LIFR
CEACAM1
PIK3R1
TRAF3IP2
JAK2
CDIP1
HLA−DQA1
CAMK2D
PAK2
RUNX1
CARD14
EGR1
FASLG
FOXO1
OTUD4
TEC
MADD
IL18BP
BRWD1
MAVS
STAT5B
BIRC5
BCL6
IFNLR1
SPATA2
CIITA
LTBR
TNFSF13
Bakay EDMD
G1
gp1
gp2
gp3
TMEM214.p1
NET39.p2
EMD.p5
FHL1.p7
LMNA.p10
SUN1.p3
FHL1.p6
FHL1.p8
SYNE1.p4
LMNA.p9
B
Cytokine signaling GO:0019221 (DE + FDR filtered)
2
1
0
−1
−2
HLA−H
TNFRSF6B
SPHK1
XAF1
IL20RB
IL23A
IL26
LCP1
CXCL2
CXCL3
IFI30
CXCL1
CXCL6
MMP3
LBP
CXCL5
IL13RA2
CXCL8
IL1B
LEP
AGPAT1
IL33
CD70
IFITM1
GREM2
HGF
IL18R1
IFI27
IRAK3
PPARG
CASP1
OAS2
IRF6
IRF7
CAMK2B
RORC
PRKN
SOX2
CEBPA
CNTF
IL17C
FCGR1A
S1PR1
CTF1
CD4
TRIM26
IL6R
STAT5A
BCL2
HSPA1B
CD80
MID1
IL7
CISH
B2M
IL15
UGCG
SHC1
ITGB1
PRKCD
TNFRSF12A
RIPK2
CIB1
CNN2
BAD
BIRC2
NR1H2
UBC
IRF3
PSMD5
RBCK1
GSTO1
P4HB
TALDO1
STX3
IKBKG
SPPL2A
ANXA2
FSCN1
IRF9
IL10RB
IRAK4
FADD
IFNGR2
RAP1B
TMSB4X
EPO
IL1RL1
FOS
MAP3K8
SOCS2
SOS1
RORA
PIK3CB
LIFR
CEACAM1
PIK3R1
TRAF3IP2
JAK2
CDIP1
HLA−DQA1
CAMK2D
PAK2
RUNX1
CARD14
EGR1
FASLG
FOXO1
OTUD4
TEC
MADD
IL18BP
BRWD1
MAVS
STAT5B
BIRC5
BCL6
IFNLR1
SPATA2
CIITA
LTBR
TNFSF13
Bakay EDMD
G1
gp1
gp2
gp3
TMEM214.p1
NET39.p2
EMD.p5
FHL1.p7
LMNA.p10
SUN1.p3
FHL1.p6
FHL1.p8
SYNE1.p4
LMNA.p9

Figure S7 (continued)

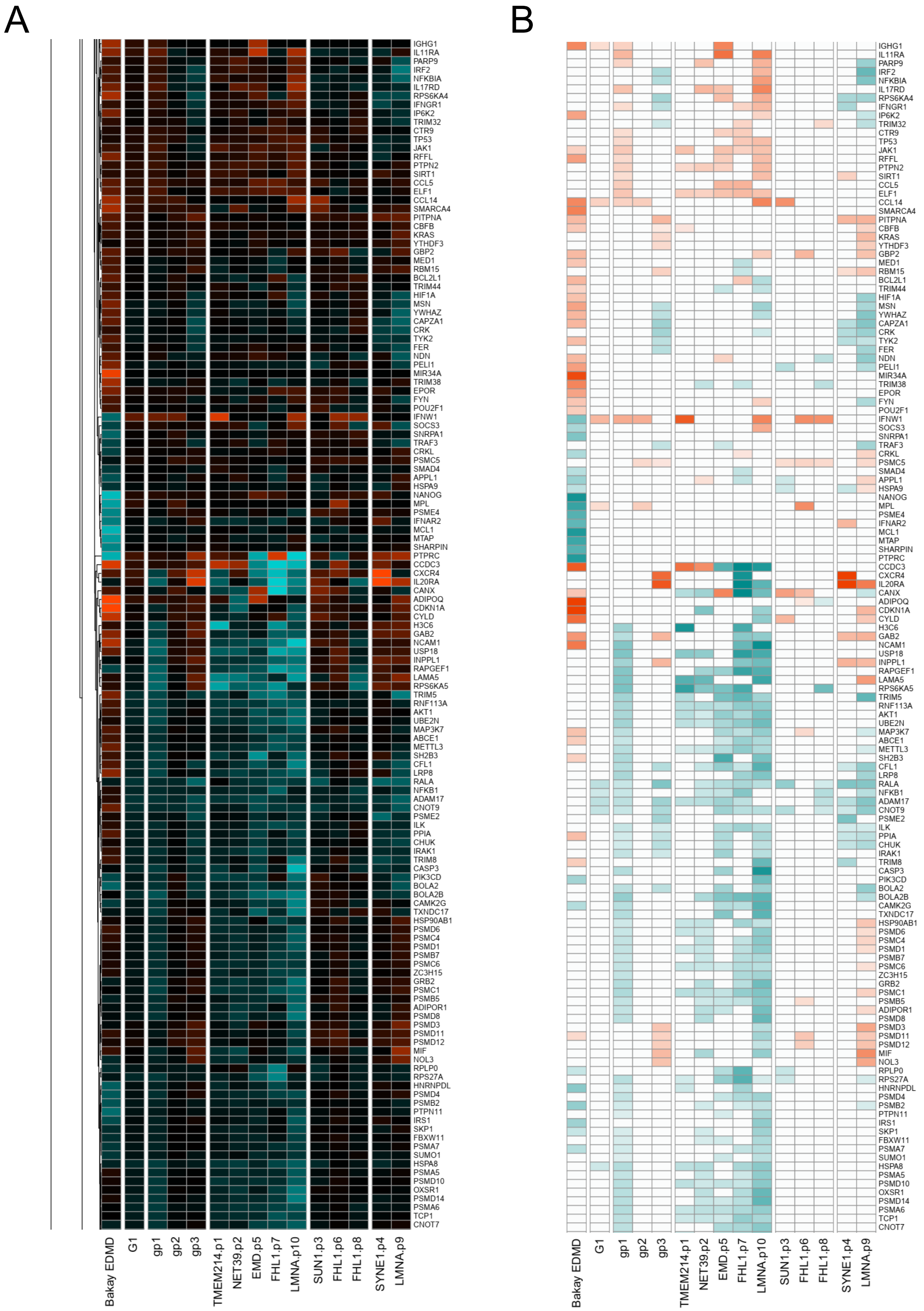

Figure S7 (continued)

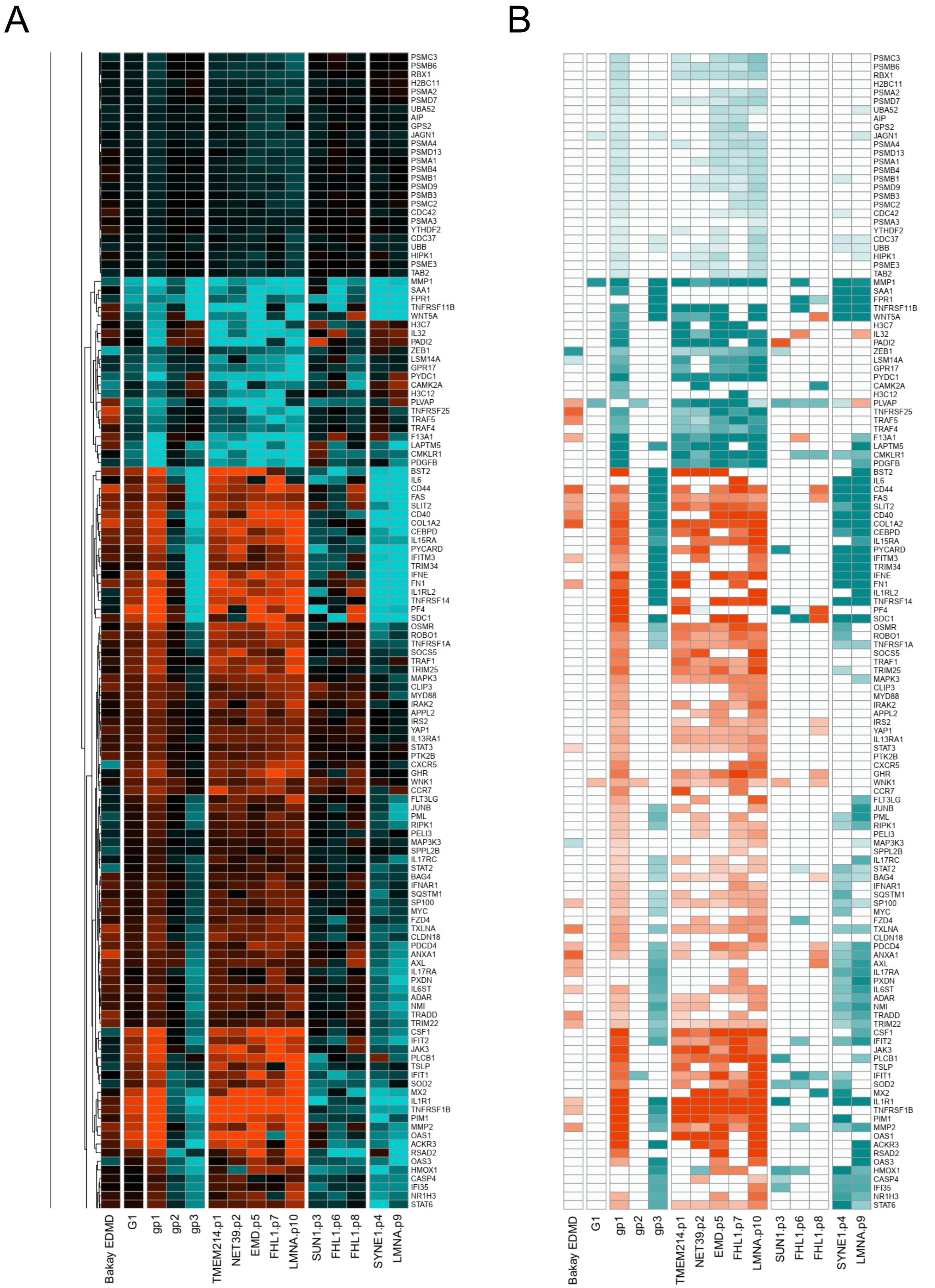

Figure S7 (continued)

A

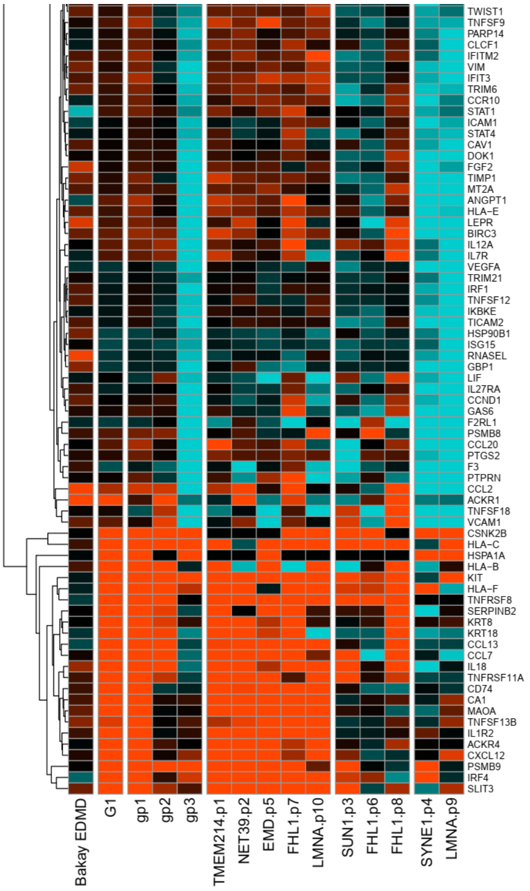


B

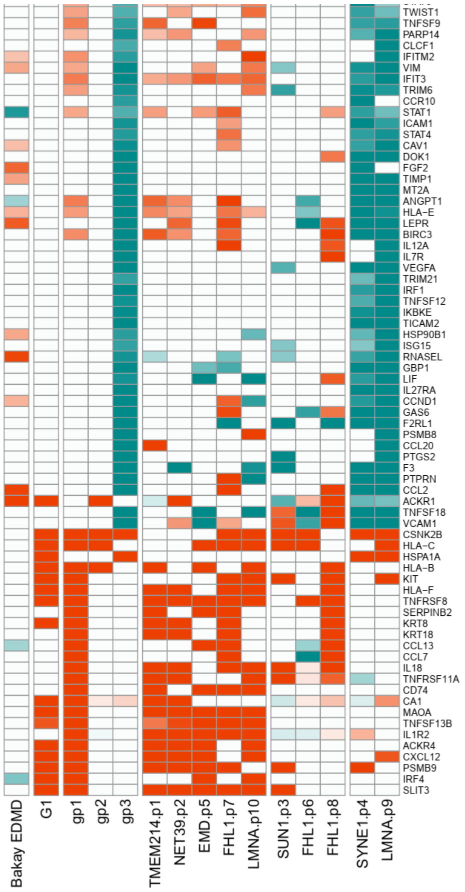

Figure S8

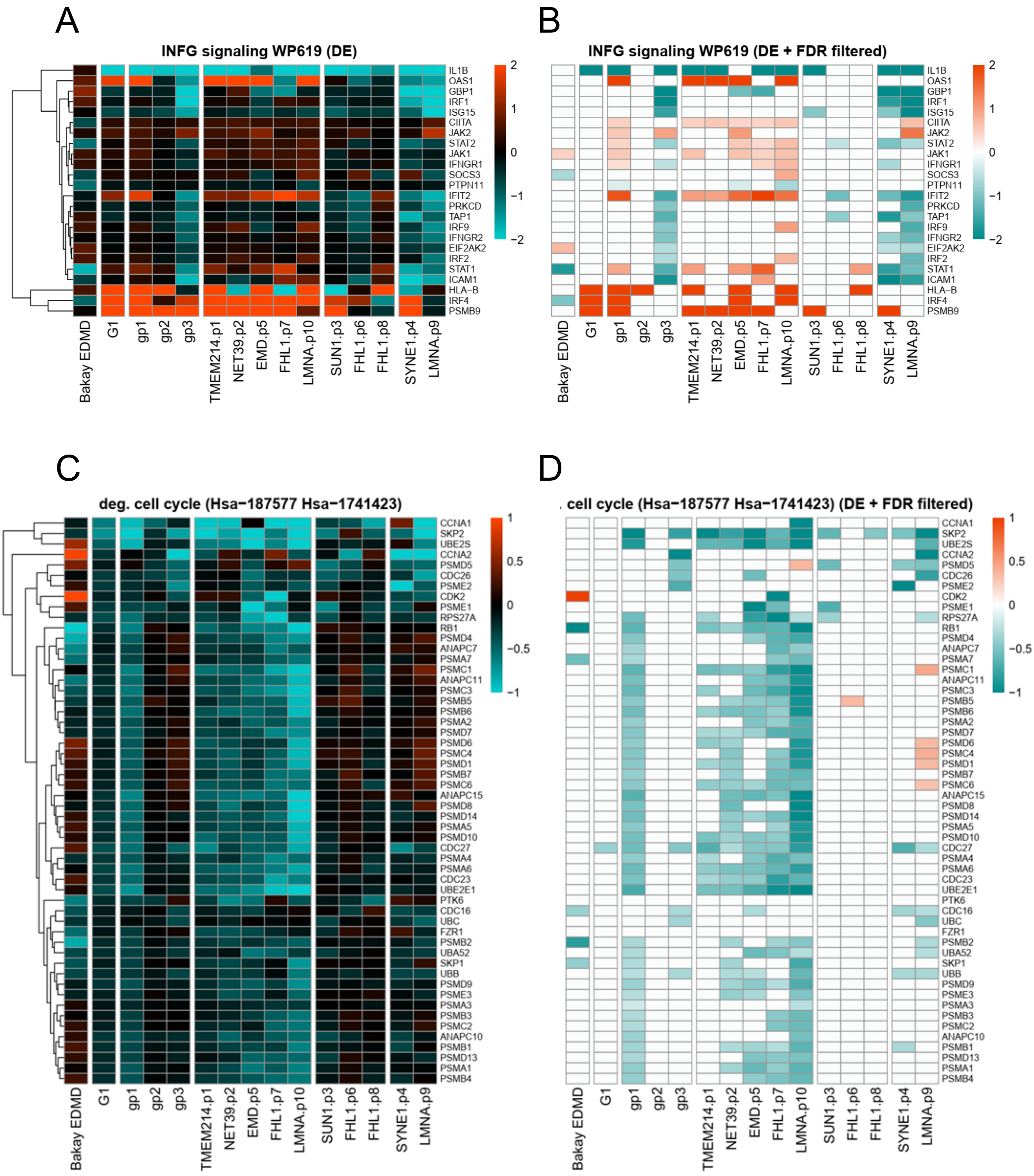
A
INFG signaling WP619 (DE)
B
INFG signaling WP619 (DE + FDR filtered)
IL1B
OAS1
GBP1
IRF1
ISG15
CIITA
JAK2
STAT2
JAK1
IFNGR1
SOCS3
PTPN11
IFIT2
PRKCD
TAP1
IRF9
IFNGR2
EIF2AK2
IRF2
STAT1
ICAM1
HLA-B
IRF4
PSMB9
Bakay EDMD
G1
gp1
gp2
gp3
TMEM214.p1
NET39.p2
EMD.p5
FHL1.p7
LMNA.p10
SUN1.p3
FHL1.p6
FHL1.p8
SYNE1.p4
LMNA.p9
C
deg. cell cycle (Hsa−187577 Hsa−1741423)
D
cell cycle (Hsa−187577 Hsa−1741423) (DE + FDR filtered)
CCNA1
SKP2
UBE2S
CCNA2
PSMD5
CDC26
PSME2
CDK2
PSME1
RPS27A
RB1
PSMD4
ANAPC7
PSMA7
PSMC1
ANAPC11
PSMC3
PSMB5
PSMB6
PSMA2
PSMD7
PSMD6
PSMC4
PSMD1
PSMB7
PSMC6
ANAPC15
PSMD8
PSMD14
PSMA5
PSMD10
CDC27
PSMA4
PSMA6
CDC23
UBE2E1
PTK6
CDC16
UBC
FZR1
PSMB2
UBA52
SKP1
UBB
PSMD9
PSME3
PSMA3
PSMB3
PSMC2
ANAPC10
PSMB1
PSMD13
PSMA1
PSMB4

Figure S9

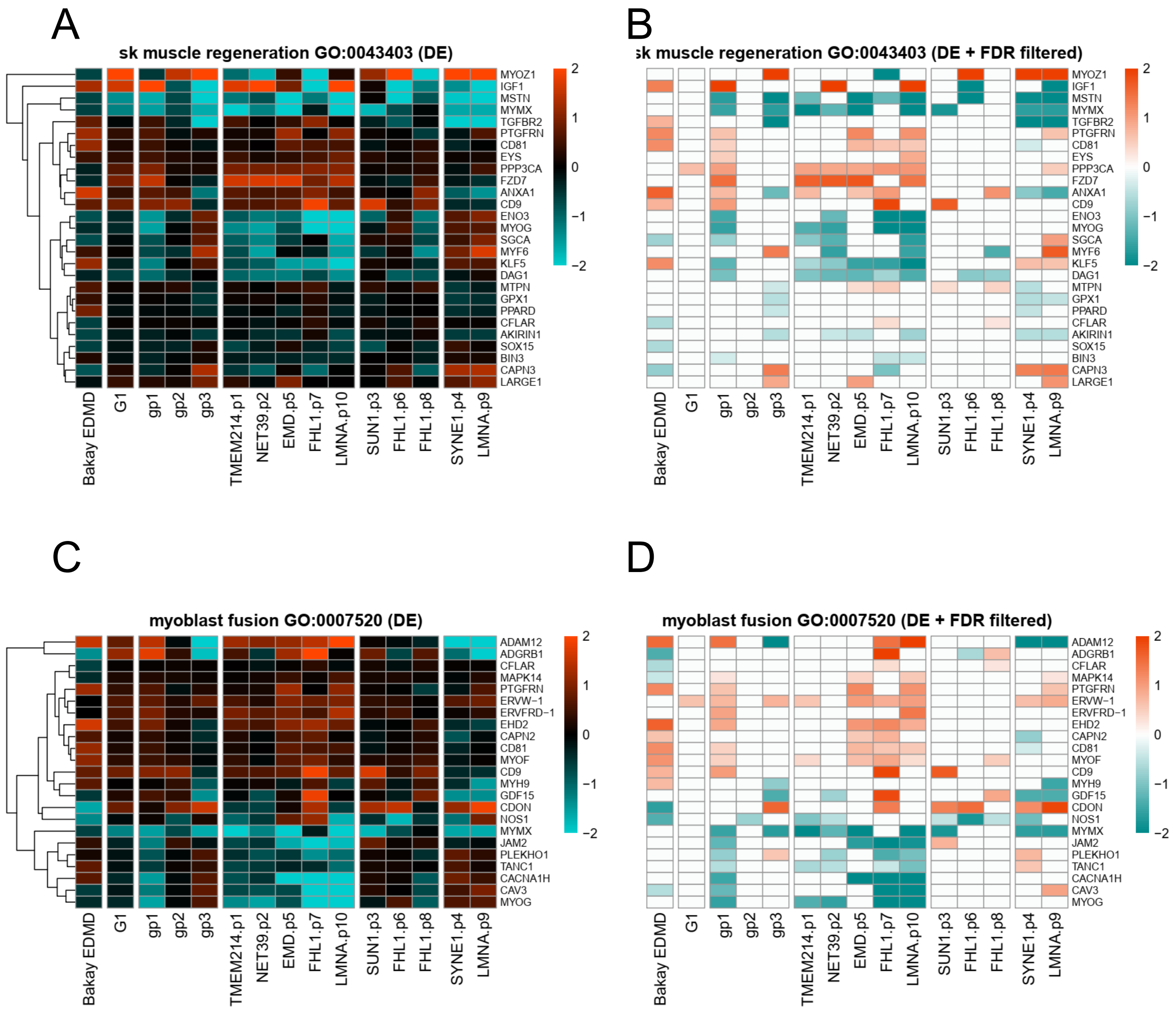

A
sk muscle regeneration GO:0043403 (DE)
B
sk muscle regeneration GO:0043403 (DE + FDR filtered)
MYOZ1
IGF1
MSTN
MYMX
TGFBR2
PTGFRN
CD81
EYS
PPP3CA
FZD7
ANXA1
CD9
ENO3
MYOG
SGCA
MYF6
KLF5
DAG1
MTPN
GPX1
PPARD
CFLAR
AKIRIN1
SOX15
BIN3
CAPN3
LARGE1
2
1
0
−1
−2
Bakay EDMD
G1
gp1
gp2
gp3
TMEM214.p1
NET39.p2
EMD.p5
FHL1.p7
LMNA.p10
SUN1.p3
FHL1.p6
FHL1.p8
SYNE1.p4
LMNA.p9
C
myoblast fusion GO:0007520 (DE)
D
myoblast fusion GO:0007520 (DE + FDR filtered)
ADAM12
ADGRB1
CFLAR
MAPK14
PTGFRN
ERVW−1
ERVFRD−1
EHD2
CAPN2
CD81
MYOF
CD9
MYH9
GDF15
CDON
NOS1
MYMX
JAM2
PLEKHO1
TANC1
CACNA1H
CAV3
MYOG

Figure S10

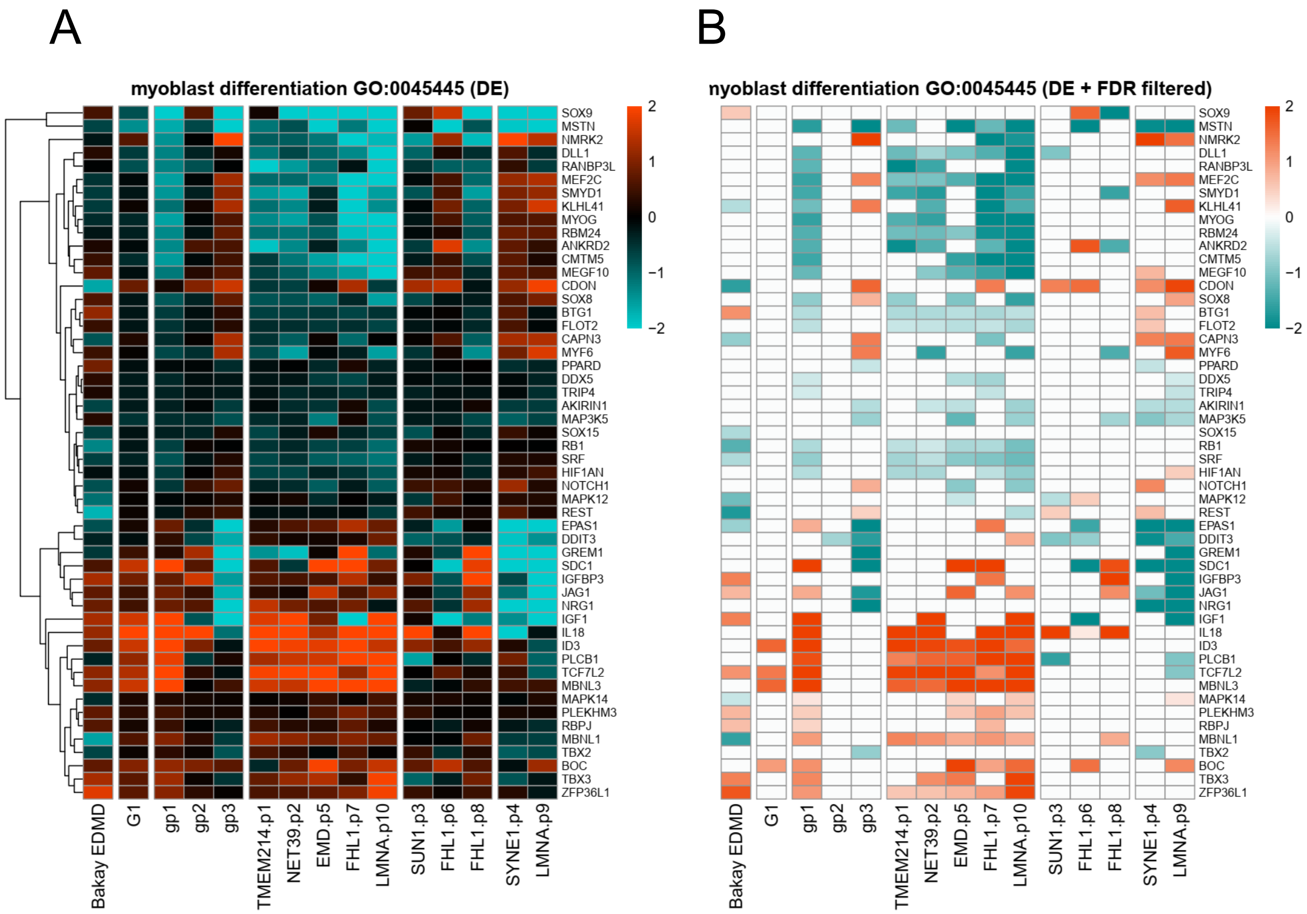
A
myoblast differentiation GO:0045445 (DE)
2
1
0
−1
−2
SOX9
MSTN
NMRK2
DLL1
RANBP3L
MEF2C
SMYD1
KLHL41
MYOG
RBM24
ANKRD2
CMTM5
MEGF10
CDON
SOX8
BTG1
FLOT2
CAPN3
MYF6
PPARD
DDX5
TRIP4
AKIRIN1
MAP3K5
SOX15
RB1
SRF
HIF1AN
NOTCH1
MAPK12
REST
EPAS1
DDIT3
GREM1
SDC1
IGFBP3
JAG1
NRG1
IGF1
IL18
ID3
PLCB1
TCF7L2
MBNL3
MAPK14
PLEKHM3
RBPJ
MBNL1
TBX2
BOC
TBX3
ZFP36L1
Bakay EDMD
G1
gp1
gp2
gp3
TMEM214.p1
NET39.p2
EMD.p5
FHL1.p7
LMNA.p10
SUN1.p3
FHL1.p6
FHL1.p8
SYNE1.p4
LMNA.p9
B
nyoblast differentiation GO:0045445 (DE + FDR filtered)
2
1
0
−1
−2
SOX9
MSTN
NMRK2
DLL1
RANBP3L
MEF2C
SMYD1
KLHL41
MYOG
RBM24
ANKRD2
CMTM5
MEGF10
CDON
SOX8
BTG1
FLOT2
CAPN3
MYF6
PPARD
DDX5
TRIP4
AKIRIN1
MAP3K5
SOX15
RB1
SRF
HIF1AN
NOTCH1
MAPK12
REST
EPAS1
DDIT3
GREM1
SDC1
IGFBP3
JAG1
NRG1
IGF1
IL18
ID3
PLCB1
TCF7L2
MBNL3
MAPK14
PLEKHM3
RBPJ
MBNL1
TBX2
BOC
TBX3
ZFP36L1
Bakay EDMD
G1
gp1
gp2
gp3
TMEM214.p1
NET39.p2
EMD.p5
FHL1.p7
LMNA.p10
SUN1.p3
FHL1.p6
FHL1.p8
SYNE1.p4
LMNA.p9

Figure S11

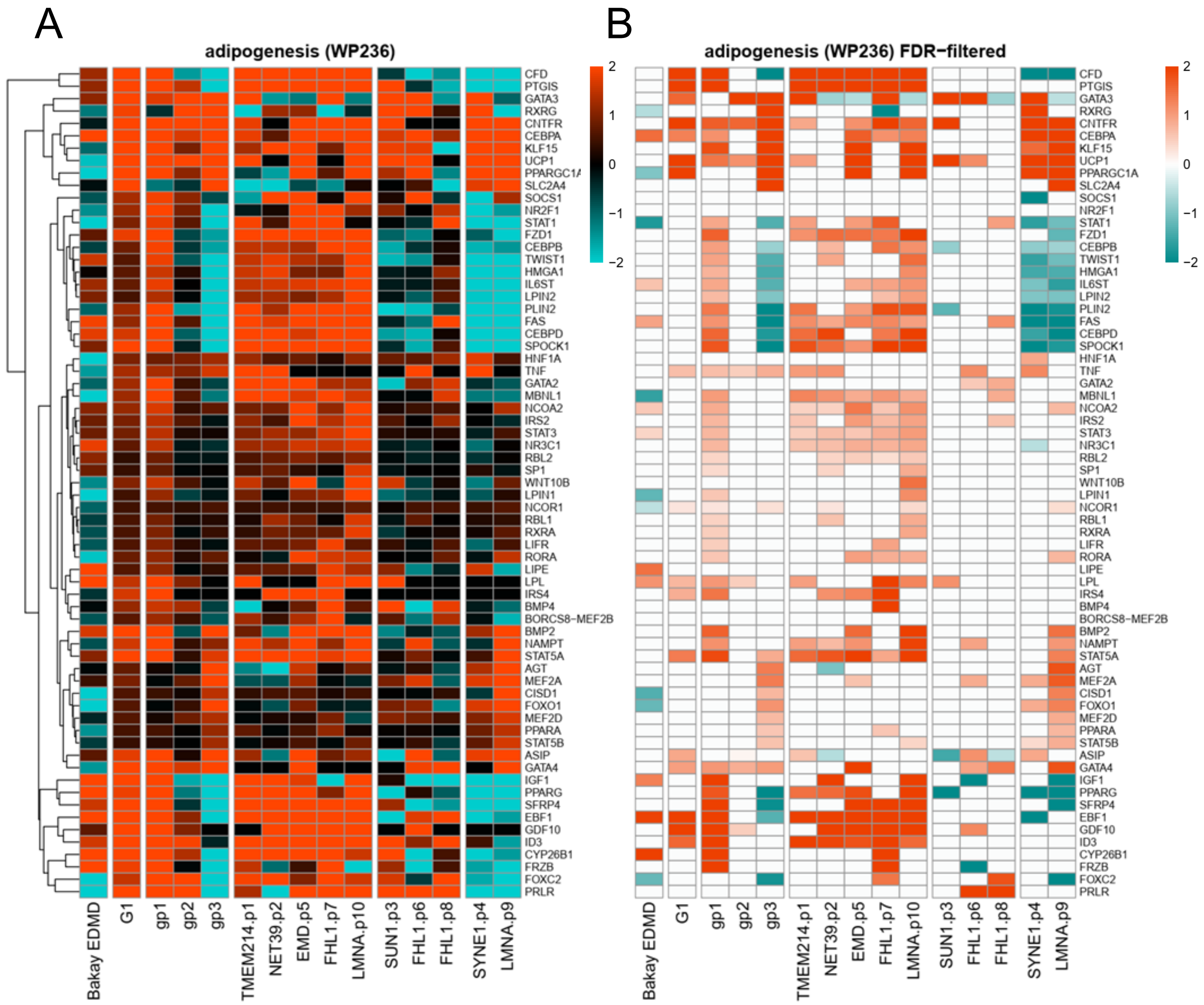
A
B
adipogenesis (WP236)
adipogenesis (WP236) FDR−filtered
2
1
0
−1
−2
CFD
PTGIS
GATA3
RXRG
CNTFR
CEBPA
KLF15
UCP1
PPARGC1A
SLC2A4
SOCS1
NR2F1
STAT1
FZD1
CEBPB
TWIST1
HMGA1
IL6ST
LPIN2
PLIN2
FAS
CEBPD
SPOCK1
HNF1A
TNF
GATA2
MBNL1
NCOA2
IRS2
STAT3
NR3C1
RBL2
SP1
WNT10B
LPIN1
NCOR1
RBL1
RXRA
LIFR
RORA
LIPE
LPL
IRS4
BMP4
BORCS8−MEF2B
BMP2
NAMPT
STAT5A
AGT
MEF2A
CISD1
FOXO1
MEF2D
PPARA
STAT5B
ASIP
GATA4
IGF1
PPARG
SFRP4
EBF1
GDF10
ID3
CYP26B1
FRZB
FOXC2
PRLR
Bakay EDMD
G1
gp1
gp2
gp3
TMEM214.p1
NET39.p2
EMD.p5
FHL1.p7
LMNA.p10
SUN1.p3
FHL1.p6
FHL1.p8
SYNE1.p4
LMNA.p9

Figure S12

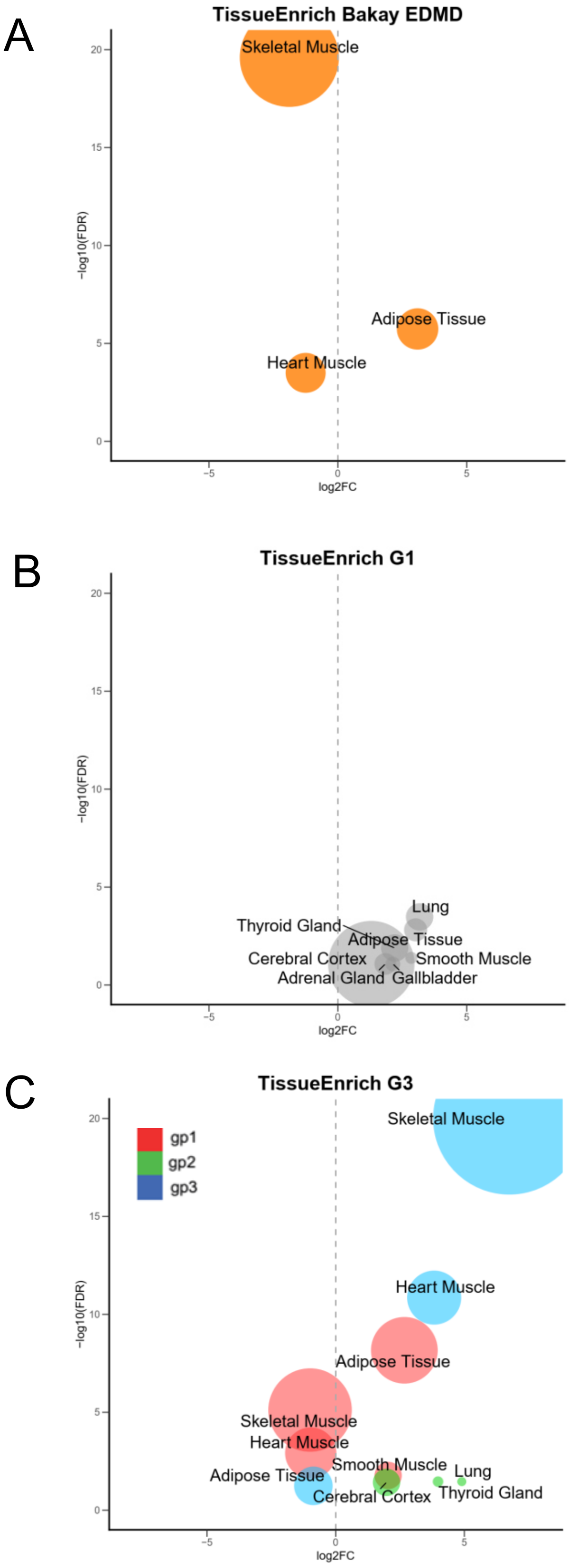

A
TissueEnrich Bakay EDMD
Skeletal Muscle
Adipose Tissue
Heart Muscle
20
15
10
5
0
−log10(FDR)
−5
0
5
log2FC
B
TissueEnrich G1
Lung
Thyroid Gland
Adipose Tissue
Cerebral Cortex
Smooth Muscle
Adrenal Gland
Gallbladder
20
15
10
5
0
−log10(FDR)
−5
0
5
log2FC
C
TissueEnrich G3
gp1
gp2
gp3
Skeletal Muscle
Heart Muscle
Adipose Tissue
Skeletal Muscle
Heart Muscle
Smooth Muscle
Lung
Adipose Tissue
Cerebral Cortex
Thyroid Gland
20
15
10
5
0
−log10(FDR)
−5
0
5
log2FC

Figure S13

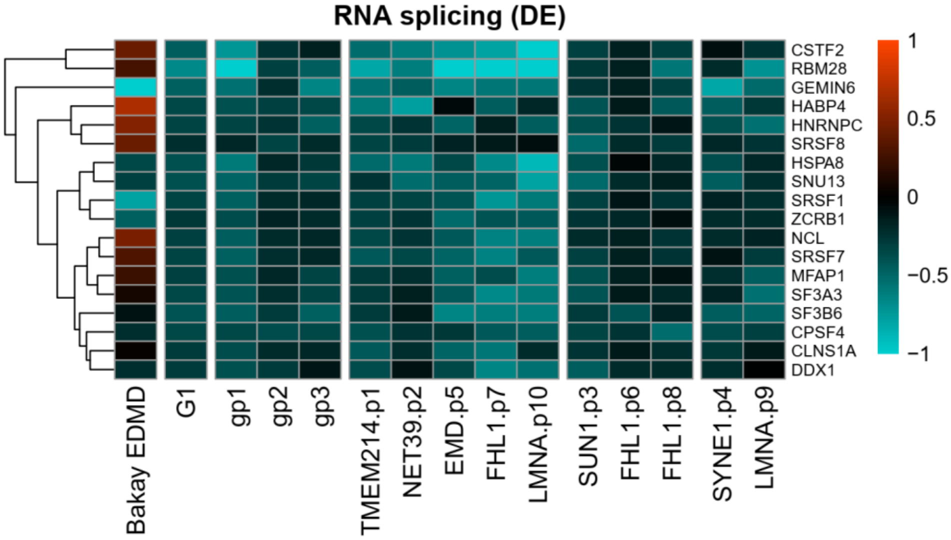
RNA splicing (DE)
CSTF2
RBM28
GEMIN6
HABP4
HNRNPC
SRSF8
HSPA8
SNU13
SRSF1
ZCRB1
NCL
SRSF7
MFAP1
SF3A3
SF3B6
CPSF4
CLNS1A
DDX1
1
0.5
0
−0.5
−1
Bakay EDMD
G1
gp1
gp2
gp3
TMEM214.p1
NET39.p2
EMD.p5
FHL1.p7
LMNA.p10
SUN1.p3
FHL1.p6
FHL1.p8
SYNE1.p4
LMNA.p9

Figure S14

A

**G1**

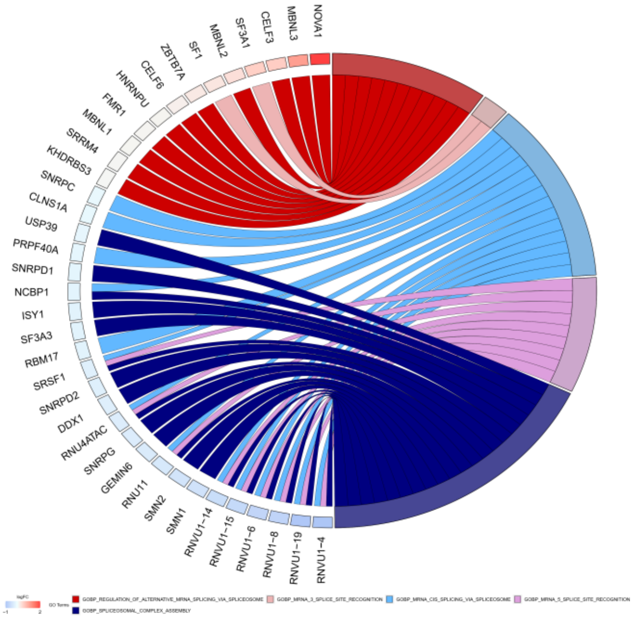


B

**gp2**

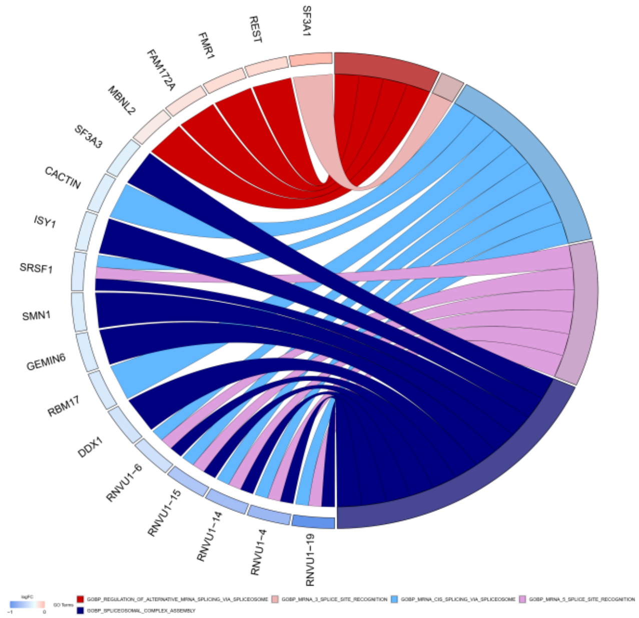


C

**gp3**

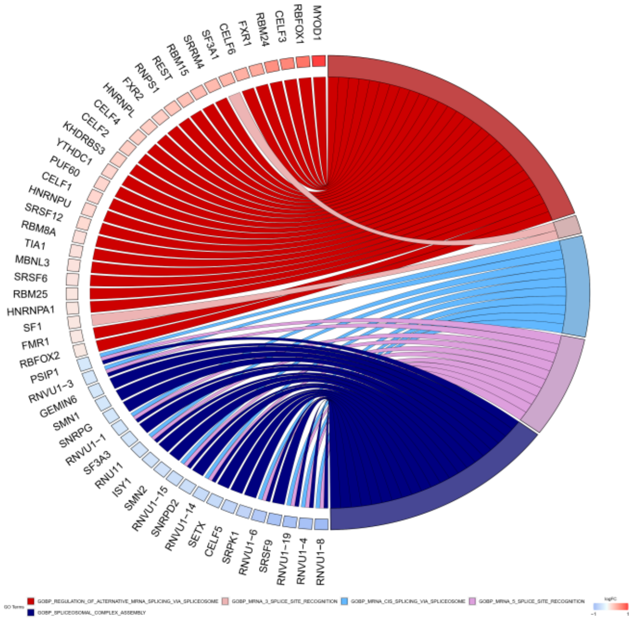

Figure S15

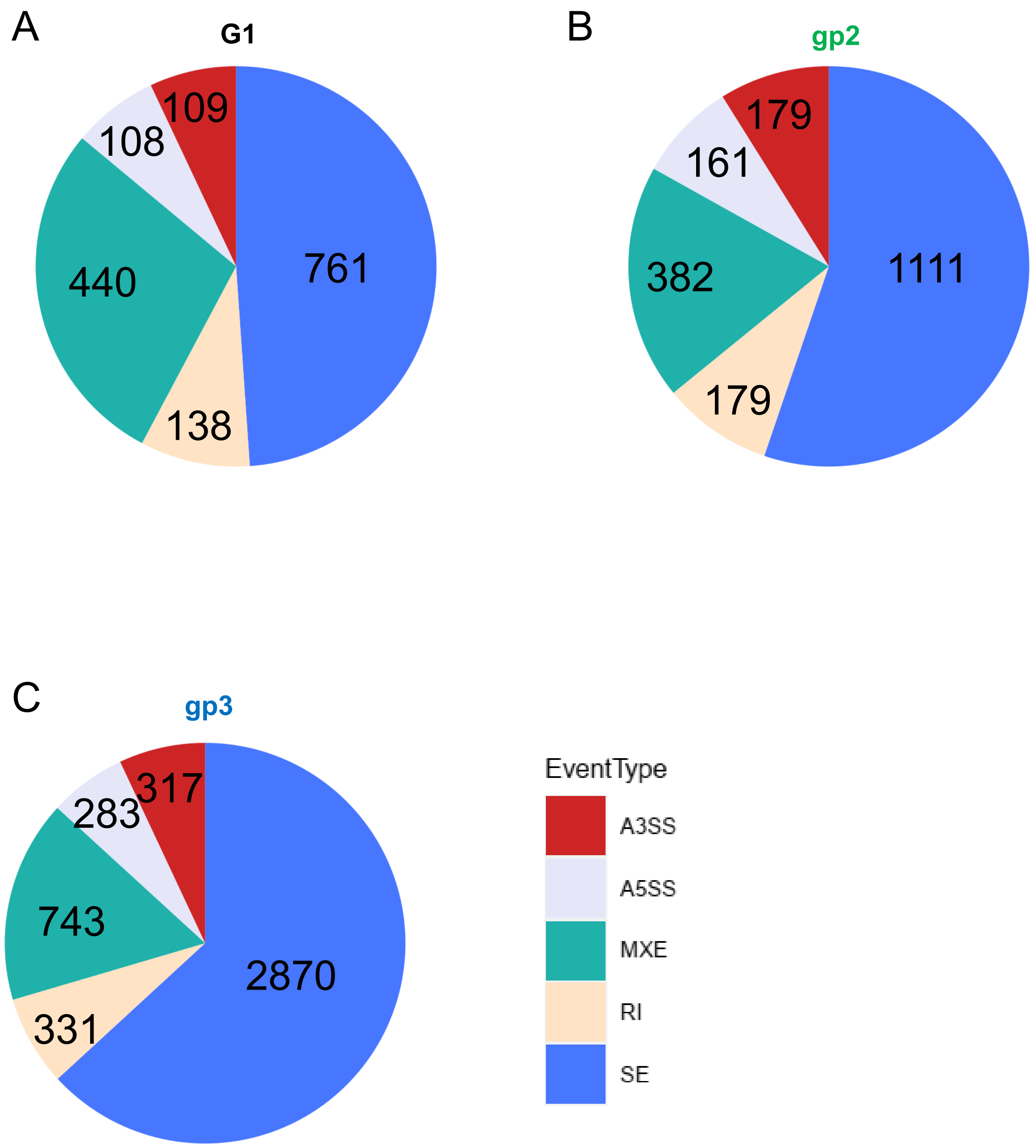
A
G1
109
108
761
440
138
B
gp2
179
161
1111
382
179
C
gp3
317
283
743
2870
331
EventType
A3SS
A5SS
MXE
RI
SE

Figure S16

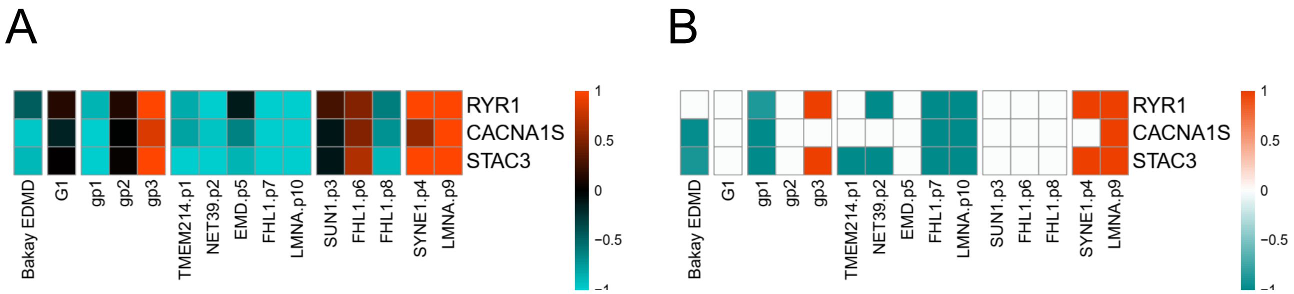
A
B
RYR1
CACNA1S
STAC3
Bakay EDMD
G1
gp1
gp2
gp3
TMEM214.p1
NET39.p2
EMD.p5
FHL1.p7
LMNA.p10
SUN1.p3
FHL1.p6
FHL1.p8
SYNE1.p4
LMNA.p9
1
0.5
0
−0.5

**Table S1. RNA quality and Sequencing depth**. RNA quality was assessed with a Bioanalyzer, and the RNA integrity number (RIN) shown for each sample where 1 = completely degraded, 10 = completely intact. A RIN value above 7 is considered ideal for RNAseq. Sequencing depth indicated by the number, in millions, of unequivocally mapped paired-end fragments (= half the number of reads) per experiment.

**Table S2. Summary of transcriptome changes and gene annotations (Hg38 genome build).** G1 (all EDMD patients together as a single group against the controls) and G10 (each individual patient analyzed separately against the controls) results provided. Positive and negative log2FC values indicate upregulation or downregulation, respectively, in EDMD.

**Table S3. Functional analysis results.** A. Bakay EDMD. g:Profiler analysis summary, representative categories. B. G1 comparison (all EDMD patients as a single group). g:Profiler analysis summary, representative categories. C. G1 comparison. GSEA analysis summary, using the C2 (pathway) geneset collection. Representative categories. D. Bakay EDMD. g:Profiler analysis, full list. E. G1 comparison (all EDMD patients as a single group). g:Profiler analysis, full list. F. G1 comparison. GSEA analysis, using the C2 (pathway) geneset collection. Full list.

**Table S4. Overlaps of misspliced genes detected by DEXSeq, rMATS and ISA.** Genes detected by all three methods shown for each of the EDMD patient subgroups as well as the G1 main group (all 10 EDMD patients analyzed together as a single group).

**Table S5. Missplicing events detected by rMATS, DEXSeq and ISA.** Full table containing all events by each method, in each of the three patient subgroups as well as the G1 main group (all 10 EDMD patients analyzed together as a single group).

**Table S6. Summary of functional categories represented in the list of 28 DE miRNAs in G1.**

**Figure S1. Enrichment of *in vitro* differentiated myotubes.** A. Human primary myotubes grown in uncoated tissue culture dishes, after 6d differentiation *in vitro*. B. Myotubes start to detach after ~30 seconds in 0.1% Trypsin solution at room temperature. C. Most myotubes detach within 2-3 minutes, with gentle swirling. D. Myoblasts may start to round up within this time frame but they remain attached to bottom of the dish. E. Supernatant removed and observed under the microscope. The sample is highly enriched for differentiated myotubes, with few contaminant myoblasts present. F. Gentle centrifugation (30 x g for 20-30 seconds) precipitates myotubes while leaving most contaminant myoblasts in the supernatant. The pellet containing the myotubes is ready to proceed to RNA extraction.

**Figure S2. Heterogeneity of gene expression changes between patients.**

A. After analysing each patient individually against the healthy controls, only 4 genes are DE in all patients at FDR 5%. B. Of the 1127 DE genes identified when analyzing all 10 patients as a single EDMD group, approximately 60% of the DE genes are altered in all 10 patients in the same direction if we disregard FDR values. C. Number of genes changing in the same direction in 1-10 patients, with and without FDR filtering, out of the 1127 DE genes identified. D. PCA analysis on patients segregates patients into three broad groups (red, green and blue). Patients with LMNA mutations contribute the most to the transcriptome profiles.

**Figure S3. t-SNE clustering analysis.** A. Individual samples, color-coded by gender. B. Individual samples, color-coded by age. C. Individual samples, color-coded by level of myotube enrichment. D. Individual samples, color-coded by patient. E. Individual samples, color-coded by condition: control or EDMD. F. Individual samples, color-coded by group: control, or subgroup 1-3.

**Figure S4. Heatmap of nuclear-encoded mitochondrial genes.** Most muscular dystrophies exhibit a downregulation of nuclear-encoded mitochondrial genes.

**Figure S5. Heatmaps showing genes differentially expressed (DE).** For each category highlighted, showing genes that are DE in at least one of Bakay EDMD, G1 (all 10 patients analyzed as one group) or subgroups gp1, gp2 or gp3. A. Collagens. B. Collagens, filtered by FDR 5 %. C. Matrix metalloproteinases. D. Matrix metalloproteinases, filtered by FDR 5 %.

**Figure S6. Heatmaps showing genes differentially expressed (DE).** For each category highlighted, showing genes that are DE in at least one of Bakay EDMD, G1 (all 10 patients analyzed as one group) or subgroups gp1, gp2 or gp3. A. Fibroatlas genes. B. Fibroatlas genes, filtered by FDR 5 %.

**Figure S7. Heatmaps showing genes differentially expressed (DE).** For each category highlighted, showing genes that are DE in at least one of Bakay EDMD, G1 (all 10 patients analyzed as one group) or subgroups gp1, gp2 or gp3. A. Cytokine signaling (GO-BP, GO:0019221). B. Cytokine signaling, filtered by FDR 5 %.

**Figure S8. Heatmaps showing genes differentially expressed (DE).** For each category highlighted, showing genes that are DE in at least one of Bakay EDMD, G1 (all 10 patients analyzed as one group) or subgroups gp1, gp2 or gp3. A. Interferon gamma signaling (WikiPathways, WP619). B. Interferon gamma signaling, filtered by FDR 5 %. C. Degradation of cell cycle proteins (Reactome, Hsa-187577, SCF/SKP2-mediated degradation of p27/p21; Reactome, Hsa-1741423, APC/C-mediated degradation of cell cycle proteins). D. Degradation of cell cycle proteins, filtered by FDR 5 %.

**Figure S9. Heatmaps showing genes differentially expressed (DE).** For each category highlighted, showing genes that are DE in at least one of Bakay EDMD, G1 (all 10 patients analyzed as one group) or subgroups gp1, gp2 or gp3. A. Skeletal muscle regeneration (GO-BP, GO:0043403). B. Skeletal muscle regeneration, filtered by FDR 5 %. C. Myoblast fusion (GO-BP, GO:0007520). D. Myoblast fusion, filtered by FDR 5 %.#

**Figure S10. Heatmaps showing genes differentially expressed (DE).** For each category highlighted, showing genes that are DE in at least one of Bakay EDMD, G1 (all 10 patients analyzed as one group) or subgroups gp1, gp2 or gp3. A. Myoblast differentiation (GO-BP, GO:0045445). B. Myoblast differentiation, filtered by FDR 5 %.

**Figure S11. Heatmaps showing genes differentially expressed (DE).** For each category highlighted, showing genes that are DE in at least one of Bakay EDMD, G1 (all 10 patients analyzed as one group) or subgroups gp1, gp2 or gp3. A. Adipogenesis (WikiPathways, WP236). B. Adipogenesis, filtered by FDR 5 %.

**Figure S12. Tissue specific gene enrichment in EDMD**. A. Using the TissueEnrich tool, the EDMD patients studied by Bakay and colleagues show enrichment of adipose tissue genes while skeletal muscle and cardiac muscle genes are downregulated. B. Considering all 10 patients as a single group (G1), adipose tissue is enriched as well as a number of other unrelated tissues although with higher p-values. C. When we analyzed each subgroup individually, group 1 exhibited the same pattern as Bakay's EDMD, group 3 showed the opposite pattern with upregulation of muscle genes and downregulation of fat genes, while group 2 was completely distinct.

**Figure S13. Heatmap of splicing genes.** Genes belonging to the RNA splicing category (GO-BP, GO:0008380) differentially expressed in G1 (all 10 patients analyzed as one group).

**Figure S14. Misregulated splicing factors**. GOchord plot for misregulated splicing factors indicates the primary change is upregulation of alternative splicing and downregulation of constitutive splicing. DE genes > |0.15| A. G1 (all 10 patients analyzed as a single group). B. Subgroup 2. C. Subgroup 3.

**Figure S15. Pie Chart of significant alternative splicing events.** Using rMATS, with psi > |0.1| in detected with rMATS. Alternative splicing (AS) events SE = exon skipping, MXE= mutually exclusive exons, RI = intron retention, A3SS = alternative 3' splice site, A5SS = alternative 5' splice site. A. AS events for G1 (all 10 EDMD patients analyzed as a single group). B. Same for subgroup 2 (gp2). C. Same for subgroup 3 (gp3).

**Figure S16. Heatmap showing expression of genes associated with malignant hyperthermia.** A. Expression relative to controls for RYR1, CACNA1S and STAC3 B. Same as A, filtered by FDR 5 %.
